# SiHDA9 interacts with SiHAT3.1 and SiHDA19 to repress dehydration responses through H3K9 deacetylation in foxtail millet

**DOI:** 10.1101/2023.02.16.528817

**Authors:** Verandra Kumar, Babita Singh, Namisha Sharma, Mehanathan Muthamilarasan, Samir V. Sawant, Manoj Prasad

**Affiliations:** National Institute of Plant Genome Research, Aruna Asaf Ali Marg, New Delhi 110067, India; Plant Molecular Biology and Biotechnology Division, National Botanical Research Institute, Rana Pratap Marg, Lucknow 226001, Uttar Pradesh, India; Department of Plant Sciences, School of Life Sciences, University of Hyderabad, Hyderabad 500046, Telangana, India

**Keywords:** Histone acetylation, H3K9ac, Histone deacetylase 9, drought, foxtail millet

## Abstract

Drought stress affects the growth and productivity of plants, where C_4_ plants can advantageously perceive and respond to the stress for their survival and reproduction. Epigenetic modifications play a prominent role in conferring drought tolerance in C_4_ plants; however, the molecular links between histone modifiers and their regulation are unclear. In the present study, we performed a genome-wide H3K9ac enrichment in foxtail millet (*Setaria italica*) and analyzed their role in regulating the expression of drought-responsive genes. The presence of H3K9ac on these genes were identified through the comparative analysis of dehydration tolerant (cv. IC4) and sensitive (IC41) cultivars of foxtail millet. A histone deacetylase, *SiHDA9*, showed significant upregulation in the sensitive cultivar during dehydration stress. *SiHDA9* overexpression in *Arabidopsis thaliana* conferred higher sensitivity to dehydration/drought stress than WT plants. We found that SiHDA9 physically interacts with SiHAT3.1 and SiHDA19. This complex is recruited through the SiHAT3.1 recognition sequence on the upstream of drought-responsive genes (*SiRAB18, SiRAP2.4, SiP5CS2, SiRD22, SiPIP1;4*, and *SiLHCB2.3*) to decrease H3K9 acetylation levels. The modulations in H3K9ac levels cause repression of gene expression and induce drought-sensitivity in the sensitive cultivar. Overall, the study provides mechanistic insights into SiHDA9-mediated regulation of drought stress response in foxtail millet.

## Main

Drought challenges the growth and productivity of crop plants^1–3^. In response, plants deploy different morpho-physiological, biochemical, and molecular responses to circumvent drought. Various histone modifications, including methylation and acetylation^1^, play a crucial role in providing tolerance against drought stress. These modifications suppress or activate the expression of several cellular reprogramming-related genes or transcription factors (TFs)^4,5^. Histone acetylation is one of the well-studied histone modifications. The enrichment or depletion of acetylation at different lysine residues of histones, including H3K9, H3K14, H3K27, and H4K5, can affect the gene expression^2,4^. The acetylation of lysine residue 9 of histone H3 (H3K9ac) is a well-studied epigenetic mark in plants. The hyper- and hypoacetylation of H3K9 results in the activation and repression of target genes, respectively. Hyperacetylation of H3K9 on the promoter region of stress-responsive genes, namely *RD29A, RD29B, RD20, ERF53*, and *RAB18*, activates their expression during drought stress-conditions^6–9^. In contrast, hypoacetylation represses their expression during drought recovery^6–9^. Hypoacetylation also suppresses the expression of negative regulatory genes, including *WRKY33*, during drought stress^6–9^. Studies also demonstrated the importance of histone acetylation (H3K9ac) under drought conditions in *Physcomitrella patens* and *Populus trichocarpa* ^5,10^.

Histone acetylation is dynamically regulated by histone acetyltransferases (HATs) and histone deacetylases (HDACs). HATs, such as GCN5, mediated H3K9ac to confer drought tolerance in Populus^5^. Conversely, HDACs, including HDA6, HDA9, and HDA19 (HD1), are also known to regulate drought tolerance mechanisms via H3K9 deacetylation^11–13^. HDA9 is a class I histone deacetylase that regulates several biological processes, such as flower development, stress responses, ABA (abscisic acid) catabolism, senescence, etc.^2,14–18^. To regulate these processes, HDA9 represses its target genes by deacetylation of H3K9/14/27ac lysine residues in coordination with different proteins such as POWERDRESS (PWR), ELONGATED HYPOCOTYL 5 (HY5), EARLY FLOWERING3 (ELF3), HIGH EXPRESSION OF OSMOTICALLY RESPONSIVE GENES 15 (HOS15), WRKY53, ABA INSENSITIVE 4 (ABI4) and HISTONE DEACETYLASE 19 (HDA19) ^12,15,16,19–25^. It has also been reported that mutation in the HDA9 resulted in stress-tolerance in Arabidopsis. These reports suggest that HDA9 is an effective epigenetic regulator of histone acetylation, which controls genes encoding for developmental and stress-responsive pathways. However, HDA9 might act as a negative epigenetic regulator of stress tolerance in plants in contrast to other HDACs that positively affect stress responses. But its precise regulatory mechanism in drought tolerance remains elusive.

Foxtail millet (*Setaria italica*) is one of the C_4_ panicoid crops known for its climate-resilient traits, including drought stress tolerance^26,27–31^. This has made foxtail millet an attractive model to study the genetic determinants underlying drought stress tolerance^30,32–34^; however, a few studies have attempted to dissect the epigenetic regulation of drought tolerance^35,36^. Besides, no systematic and detailed investigations have been made to decipher the role of epigenetics in regulating the genes involved in drought stress responses.

In the present study, we analyze the genome-wide H3K9ac distribution and gene expression in dehydration-tolerant ‘IC-403579’ (IC4) and sensitive ‘IC-480117’ (IC41) cultivars of foxtail millet in response to dehydration stress^27,37,38^. The GO analysis showed that the chromatin organization was enriched in the up-regulated, whereas glucosyltransferase activity and transcription factors (TFs) pathways were enriched in down-regulated DEGs in IC41 respective to IC4 cultivar post-24 h of dehydration stress. The genes encoding for histone modifiers that play roles in chromatin organization were significantly up-regulated in IC41. These genes include *SiHDA9 (Histone deacetylase 9) SiHAC2 (Histone acetyltransferase of the CBP family 2), SiHAT3.1 (Homeodomain box protein), SiPIE1 (Photoperiod-independent early flowering 1*) and *SiBRM1* (*BRAHMA1*). *In-silico* analysis of protein-protein interaction showed that these proteins can form a multiprotein complex with SiHDA9. Overexpression of SiHDA9 in *Arabidopsis thaliana* (Col-0) resulted in higher sensitivity to drought stress in transgenic lines. Further analysis of SiHDA9 showed its interaction with SiHAT3.1 and SiHDA19. This trimeric protein complex (SiHDA9-SiHAT3.1-SiHDA19) could binds on the promoter of drought-responsive genes, including *SiRAB18, SiRAP2.4, SiP5CS2, SiRD22, SiPIP1;4* and *SiLHCB2.3* through SiHAT3.1 recognising sequence (T(A/G)(A/C)ACCA). This decreases their H3K9ac levels leading to gene repression, which impacts the drought tolerance in the sensitive cultivar (IC41) compared to the tolerant cultivar (IC4). Overall, this study delineates the importance of histone acetylation of H3K9 for drought tolerance and provides mechanistic insights into the role of SiHDA9 in removing the acetylation to negatively regulate the drought-tolerance mechanism.

## Results

### The significant perturbation in the dehydration-responsive gene network in both dehydration-tolerant and sensitive cultivars of foxtail millet

Previously, foxtail millet cultivars IC4 (dehydration-tolerant) and IC41 (dehydration-sensitive) were reported to differ contrastingly in their tolerance to PEG-induced dehydration stress^32,37,38^. Therefore, the transcriptomes of these two cultivars at 0 h and 24 h post-PEG treatments were sequenced to identify the differentially expressed genes and their associated co-expression networks (Supplementary Table 1)^28^. The analysis identified a total of 7097 (3625 up and 3472 down) and 7712 (4594 up and 3118 down) differentially expressed genes (DEGs; >1 Log2 fold change, FDR<0.05) between 24 h and control (0 h) samples in IC4 and IC41, respectively (Fig. 1a,b, Supplementary Table 2). The DEGs of IC4 and IC41 were compared and categorized into G1-G8 groups based on their expression pattern (Fig. 1c, Supplementary Table 3). All groups showed significant differences in expression levels between the cultivars (Fig. 1d, Extended data Fig. 1,2). Pathway analysis of all the groups revealed enrichment of DEGs encoding for developmental and stress-responsive pathways (Fig. 1e, Supplementary Table 4). These pathways include cellular and metabolic processes (enriched in G1-6) in the case of the development, whereas response to stimulus and chemical (enriched in G1-6 and G8) in the stress-response category (Fig. 1e). Interestingly, the genes involved in pathway, response to water, were enriched only in the group G3 (unique IC4-UP; tolerant cultivar) with G1 and G2 (common genes), whereas genes involved in chromatin organization, chromosome organization, negative regulation of the biological process and response to heat pathways were enriched in the G5 (unique IC41-UP; sensitive cultivar) with G1 group of DEGs. The photosystem (I and II) and antioxidant activity-related pathways encoding genes belonged to G6 (unique IC41-DN) with G2. In contrast, water transmembrane transporter activity and water channel activity-related pathway encoding genes were enriched only in G6. The regulation of response to stimulus, regulation of stress response and regulation of stomatal movement pathway encoding genes were found exclusively in group G5. These results suggest that the pathways underlying dehydration stress tolerance are more prominently activated in IC4 than in IC41. On the contrary, the chromatin organization pathway encoding genes were more activated in the sensitive cultivar (IC41).

**Fig. 1.**
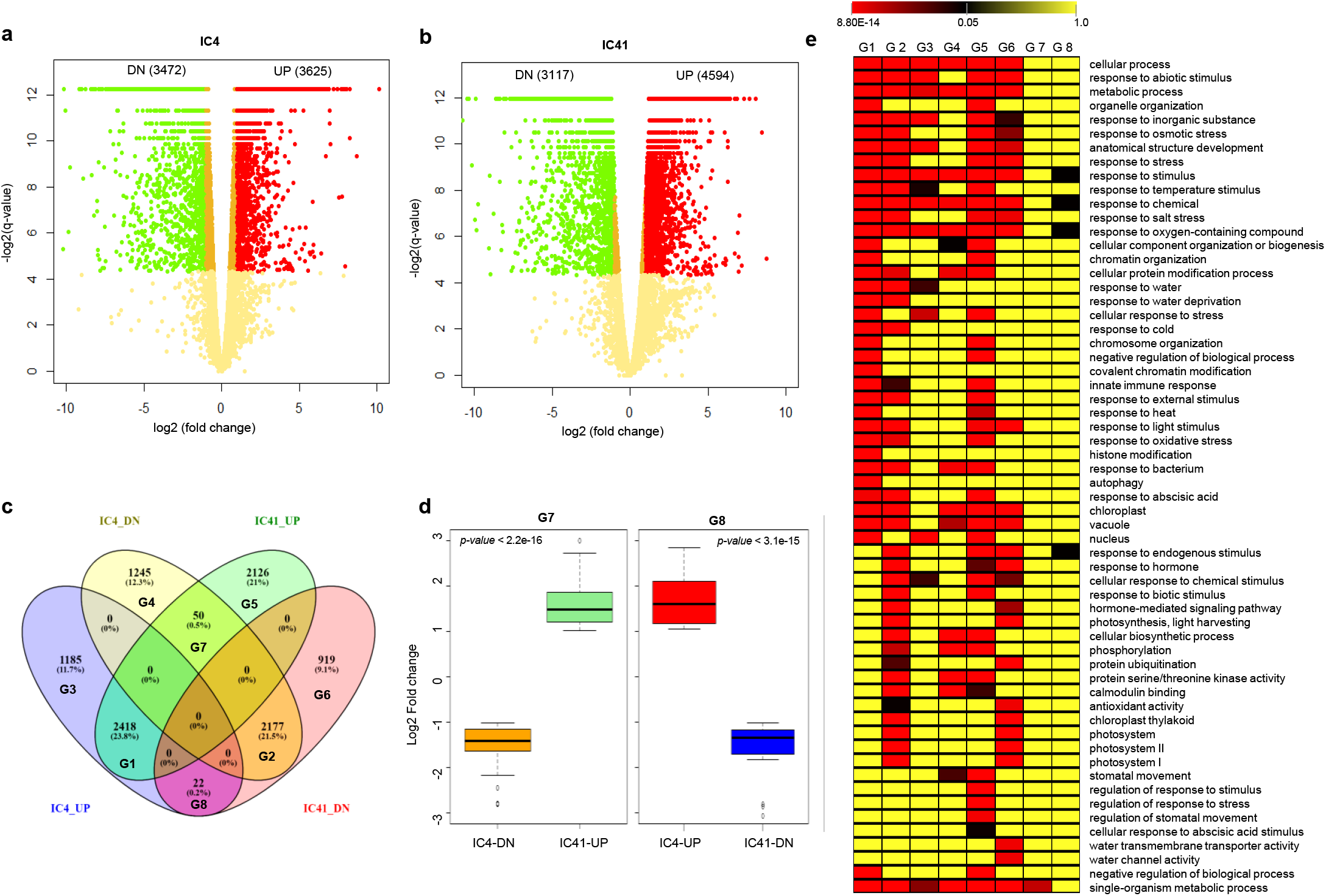
Several differentially expressed genes (DEGs) post 24hr PEG treatment were encoding to the stress-related pathways in foxtail millet. **a & b)** Volcano plots showing the significant DEGs (>1 Log2 Fold Change, FDR<0.05) after 24hr PEG treatment in tolerant (IC4) and sensitive (IC41) cultivars of foxtail millet. The red and green color dots represent the UP and Down (DN)-regulated genes, respectively, whereas the yellow dots show genes with no significant differences. **c)** Venn diagram showing the common UP/DN (similar expression pattern), unique UP/DN and genes having opposite expression patterns between the cultivars. **d)** Box plot showing the difference in the genes with opposite expression in both cultivar. **e)** Heat map showing the pathways enrichment in the different categories of genes. The G1-8 showing to the groups of the genes in the different categories such as common with similar expression pattern G1 IC4_IC41-UP (2418 genes) and G2 IC4_IC41-DN (2177 genes), with unique expression pattern G3 IC4_UP (1185 genes), G4 IC4_DN (1245 genes), G5 IC41_UP (2126 genes) and G6 IC41_DN (919 genes) and common with opposite expression pattern G7 IC4-DN_IC41-UP (50 genes) and G8 IC4-UP_IC41-DN (22 genes).

### Chromatin organization pathway encoding genes were more enriched in the dehydration-sensitive cultivar compared to the tolerant cultivar

To better understand the regulatory networks involved in dehydration tolerance, the up- and down-regulated genes were identified by calculating the Log2 FC ratio (IC41/IC4) between the cultivars. The pathway analysis (Parametric Analysis of Gene Set Enrichment) by AgriGO (v2) online tool identified several enriched pathways in IC41 in comparison to IC4 (Fig. 2a, Supplementary Table 5). These pathways include chromatin organization, covalent chromatin modification and negative regulation of gene expression. In contrast, genes involved in photosynthesis, response to cold, aging, cell wall organization or biogenesis, leaf senescence, response to ethylene, water channel activity, and peroxidase activity pathways were included in the down-regulated dataset of IC41. Other pathways, such as response to the hormone, response to temperature stimulus, response to radiation and response to light stimulus, were also enriched in down-regulated dataset in IC41 compared to IC4.

**Fig. 2.**
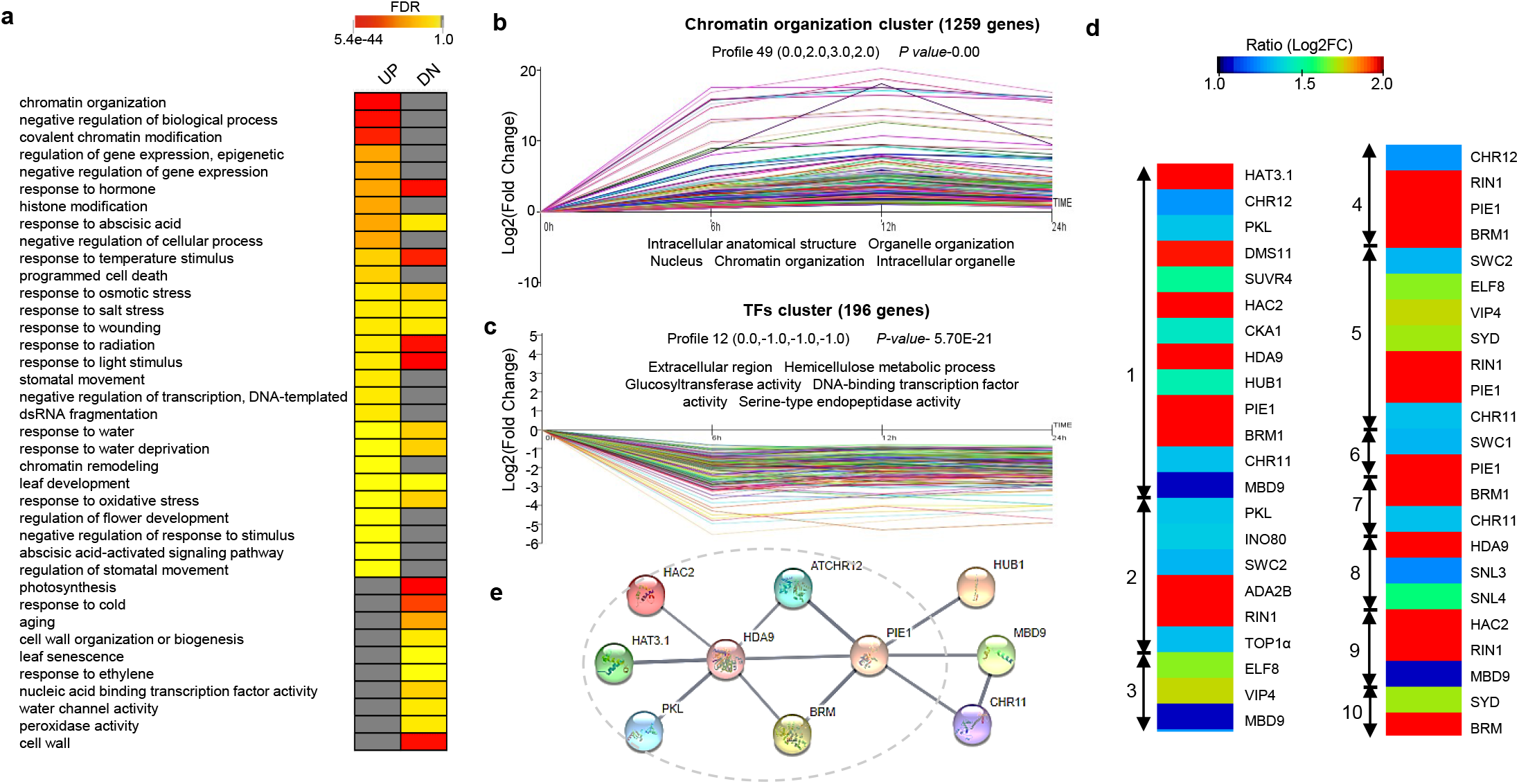
Chromatin organization (enriched) and responses to water deprivation (depleted)-related pathways modulated in the IC41 respective to the tolerant cultivar in foxtail millet. **a)** Heat map showing the affected pathways in the IC41 (UP- and Down-regulated genes) respective to the IC4. **b, c)** A cluster analysis revealed two main clusters in the PEG treatment time-course experiment in IC41 respective to the IC4 cultivar. The chromatin organization cluster having **(b)** 1259 genes; reflects the majority of dehydration-induced genes and the TFs cluster **(c)** 196 genes, showed the dehydration-repressed genes. Clusters visualize the log2 (Fold change) expression dynamics after PEG-treatment. The five strongest enriched GO terms shown for each cluster. For each time point (0, 6, 12 and 24hr), the average expression of IC41 was normalized with the average expression of IC4 (IC41/IC4) of RNA-sequencing data. **d)** Heat map showing the ratio of IC41/IC4 Log2FC (fold change) in chromatin organization pathway encoding genes, UP-regulated (Log_2_FC) in the IC41 in comparison to the IC4 cultivars. The numbers showed the top 10 sub pathways; 1 Chromatin organization, 2 Chromatin remodeling, 3 Histone modification, 4 DNA duplex unwinding, 5 Flower development, 6 Histone exchange, 7 Positive regulation of cellular response to heat, 8 Histone deacetylation, 9 Histone acetylation, 10 Organ boundary specification between lateral organs and the meristem. **e)** String result of the protein-protein interacting network in chromatin organization pathway enriched after PEG treatment in the IC41 cultivar. Proteins are represented by different colored nodes while the protein-protein association is represented by grey lines or edges at a high confidence level (0.700). Grey dotted circle showing the direct SiHDA9 associated genes.

To understand the dynamics of transcriptional regulatory network during dehydration stress, gene clustering was performed using the FPKM ratio of significant DEGs at different time points, *viz*., 0, 6, 12 and 24 h after PEG-treatments. The analysis categorized DEGs into different clusters with comparable expression patterns over time to enable the visualization of multifaceted expression dynamics with over-represented functional annotations (Fig. 2b, Extended Data Fig. 3, Supplementary Table 6). The largest up-regulated cluster was the “Chromatin organization,” and enriched for GO terms related to intracellular anatomical structure, organelle organization, nucleus, chromatin organization and intracellular organelle (Fig. 2b). In contrast, the largest cluster in down-regulated genes was the “Transcription Factors (TFs) cluster,” which is associated with enriched GO terms, namely, extracellular region, hemicellulose metabolic process, glucosyltransferase activity and serine-type endopeptidase activity (Fig. 2c). Further, the chromatin organization pathway was sub-classified into sub-pathways, including chromatin remodelling, histone modification, histone acetylation, and histone deacetylation (Fig. 2d). There were several genes encoding these pathways such as *SiHDA9* (*Histone deacetylase 9*), *SiHAC2* (*Histone acetyltransferase of the CBP family 2*), *SiHAT3.1* (*Homeodomain box protein*), *SiPIE1* (*Photoperiod-independent early flowering 1*) and *SiBRM1* (*BRAHMA1*), which showed higher expression level in IC41 in comparison to IC4 (Fig. 2d). The prediction of protein-protein interactions by STRING tool showed that these proteins can form a multiprotein complex with SiHDA9 (Fig. 2e). Thus, these results indicated that SiHDA9 might play a role in repression of genes involved in stress-responsive pathways by regulating histone acetylation dynamics in the sensitive cultivar, as HDA9 is a known histone modifier that can cause histone deacetylation to repress gene expression^12,15,16,19–25^.

### Dynamics of H3K9ac control dehydration stress tolerance mechanism

Several studies demonstrated the involvement of H3K9ac in regulating dehydration/drought responses to confer tolerance^5,10^. In Arabidopsis, it has also been shown that HDA9 regulates the H3K9ac dynamics^12,20,25^. The transcriptome data in the present study also indicates the higher expression of *SiHDA9* after 24 h dehydration stress in IC41 than in IC4 cultivar. Therefore, to identify the modulated H3K9ac regions, chromatin immunoprecipitation (ChIP) was performed by using the anti-H3K9ac antibody in the 0 h (C) and 24 h PEG-treated (P) samples followed by sequencing (ChIP-Seq). The quality-filtered reads were mapped on the reference genome of *Setaria italica* (Supplementary Table 1; Extended data Fig. 4) to identify significantly enriched H3K9ac peaks in IC4-C, IC4-P, IC41-C, and IC41-P samples. The distribution (± 2Kb from the TSS; transcription start site) of these H3K9ac peaks showed a higher enrichment near the TSS (Extended data Fig. 5). Genomic distribution of these peaks showed that most of them were located in the genic regions (except IC4-C); maximum onto the exons followed by promoter and introns, and lowest to the TTS (transcription termination site) in all the samples (Extended data Fig. 5a,b). Notably, the estimation of H3K9ac level (FPKM) on the stress-responsive unique DEGs showed a higher and lower enrichment of H3K9ac on the gene with ± 1Kb or promoter (−1Kb) region post-PEG treatments (24 h) in IC4 and IC41, respectively (Extended data Fig 7 a-d). These results prompted further experiments to determine the biological significance of H3K9ac-modified sites.

For the correlation study, the ChIP-seq data was integrated with the gene expression data (RNA-seq) to identify the DEGs exhibiting differential H3K9ac levels, mainly on ± 2Kb from TSS. The analysis identified four possible combinations after the correlation of differential gene expression with differential H3K9ac levels in both cultivars. These combinations fall into two categories. The first category is positively correlated DEGs (blue dots); 1) DEGs with increased H3K9ac level, which induced gene up-regulated (cUP-gUP) and 2) DEGs with decreased H3K9ac level, which induced gene down-regulated (cDN-gDN). The second category is negatively correlated DEGs (orange dots); 3) DEGs with increased H3K9ac level, which induced gene down-regulated (cUP-gDN) and 4) DEGs with decreased H3K9ac level, which induced gene up-regulated (Fig. 3 a,b). The H3K9ac is an activation mark^2,5,18,39^; therefore, the positively correlated genes (cUP-gUP and cDN-gDN) might epitomize a direct effect on gene expression. Thus, we primarily focused on identifying the DEGs directly regulated by H3K9ac modulations. The positively correlated DEGs, cUP-gUP and cDN-gDN (317 and 451 DEGs, respectively, in IC4 and 135 and 188 DEGs, respectively in IC41) of both cultivars were used in the pathway analysis (GO enrichment) to explore their functional significance. The study showed the overrepresentation of several developmental and stress-related pathways in both combinations of DEGs in IC4 and IC41 (Fig. 3c, Supplementary Table 7). Interestingly, the stress-related pathways such as response to water and water deprivation, response to cold, response to temperature stimulus, response to osmotic stress, response to acid chemical, response to salt stress and response to abscisic acid (ABA) were overrepresented only in cUP-gUP but not in cDN-gDN gene set of IC4, whereas, these pathways significantly enriched in both the gene sets of IC41 cultivar. However, a lower significant (q-value) enrichment was found of these pathways in the cUP-gUP of IC4 than cUP-gUP of IC41 cultivars. These results indicated that H3K9ac dynamics might differently regulate diverse stress-responsive genes in IC4 and IC41. Therefore, the cumulative H3K9ac and expression level of stress-responsive genes of positively correlated genes of IC4-cUP-gUP, IC4-cDN-gDN, IC41-cUP-gUP and IC41-cDN-gDN was estimated in both the cultivars. The H3K9ac and expression level corresponds to each other in all the categories of stress-responsive genes in IC4 and IC41 (Fig. 3d-g). Strikingly, the analysis also revealed significant differences in the expression level of these categories of genes between both cultivars. Thus, the results depicted the lowered histone acetylation and expression level of stress-responsive genes in IC41. In agreement with these observations, the RNA-seq analysis also showed that the chromatin organization pathway was overrepresented in the IC41 enlists *SiHDA9* gene (Fig. 2a-d). *SiHDA9* showed an enhanced expression level in IC41 than the IC4 cultivar after 24 h PEG treatment (Fig. 2d). Thus, the overall results suggest that SiHDA9 might be decreasing the histone acetylation level to suppress the expression of stress-responsive genes, which in turn make the plants sensitive to the different stresses like dehydration/drought as in the case of IC41 (dehydration sensitive cultivar).

**Fig. 3.**
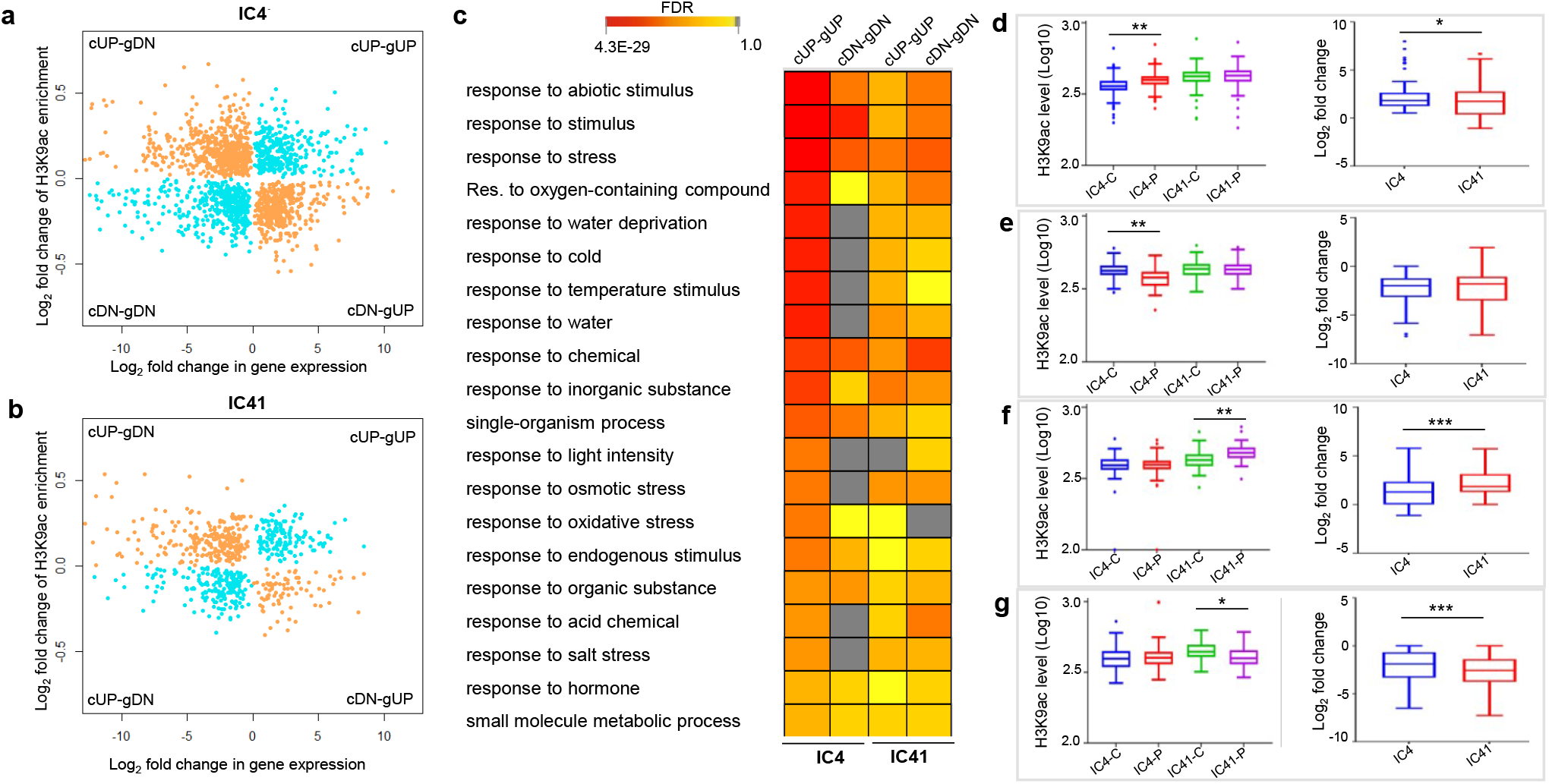
Integration of ChIP-seq and RNA-seq data identified drought stress-responsive genes associated with H3K9ac genic Regions. Scatter plots showing the correlation between gene expression (Log_2_ fold change) and H3K9ac enrichment (Log_2_ fold change) at genic regions; promoter and or gene body in **a)** IC4 and **b)** IC41. cUP- increased H3K9ac level, cDN- decreased H3K9ac level, gUP- gene up-regulation, gDN- gene down-regulation. Blue and golden colored dots represent the positively and negatively correlated genes between RNA-seq and ChIP-seq data, respectively. **c)** Heat map showing the significantly enriched pathways (FDR<0.05) in the positively correlated genes; gene expression and H3K9ac level both increased (cUP-gUP) or both decreased (cDN-gDN)) post 24hr PEG treatment in comparison to the 0hr (control). Boxplot showing the H3K9ac level (Log10 of read coverage) and expression level (Log2Fold change) of response to stress encoding genes found in **d)** IC4-cUP-gUP **e)** IC4-cDN-gDN **f)** IC41-cUP-gUP and **g)** IC41-cDN-gDN categories of correlated genes (after integration of ChIP-seq and RNA-seq data of control and PEG-treated (24hr) samples in both cultivars (IC4 and IC41). Asterisks represent the Student’s t-test: * *P* < 0.05, ** *P* < 0.01, and*** *P* <0.001.

### *SiHDA9* negatively controls dehydration/drought stress-responses

The RNA-seq and ChIP-seq data indicated that SiHDA9 might control stress responses by histone deacetylation in IC41. To corroborate this supposition, first, we overexpressed *SiHDA9* (a similar coding region in both the cultivars; Extended Data Fig. 8) in the yeast (*S. cerevisiae*). Then the *SiHDA9* overexpressing and empty vector (pYES2) containing yeast cells were exposed to dehydration stress (30% PEG) for 24 h. *SiHDA9* overexpressing yeast cells showed reduced growth than the control cells; however, no significant differences were observed between empty vector containing and *SiHDA9* overexpressing cells in unstressed conditions (Extended Data Fig. 9). These observations suggested that the SiHDA9 controlled dehydration stress-responsiveness. To validate this, *SiHDA9* was overexpressed in Arabidopsis. The transgenic lines were generated after transforming the construct, and positive lines were screened by antibiotic (Kanamycin) selection followed by PCR in T_2_-generation (Extended Data Fig. 10a,b). Additionally, the expression of *SiHDA9* was also confirmed by qRT-PCR in T_2_ lines (Extended Fig. 11a). We selected two higher *SiHDA9* expressing positive T_2_ lines (*SiHDA9-OE1* and *SiHDA9-OE2*) for further analysis in T_3_ generation. The qRT-PCR detected *SiHDA9* expression in both the lines (in T_3_ generation), and it was absent in the Col-0 (wild-type) (Fig. 4a). Next, the stress treatments were performed in the Petri-dishes containing 150mM and 300mM concentrations of mannitol with 1/2MS agar (0.8%) medium. The *SiHDA9* overexpression lines showed stunted growth compared to the Col-0 at both mannitol concentrations, which ultimately caused a significant reduction of biomass in the transgenic lines. However, no significant alterations were found between untreated (control) Col-0 and transgenic lines (Extended Fig. 11b,c).

**Fig. 4.**
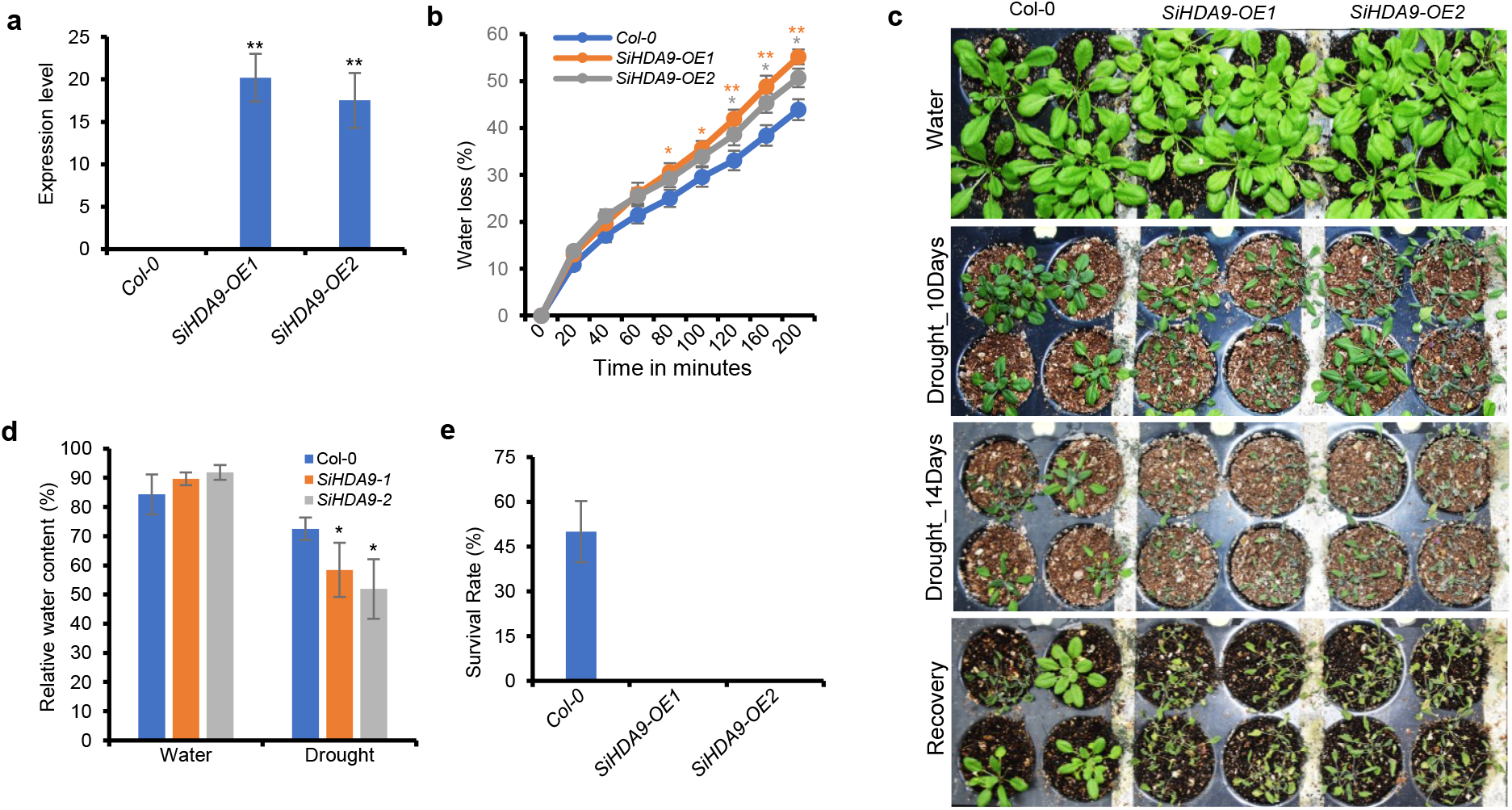
The overexpression lines of *SiHDA9* in Arabidopsis are hypersensitive to drought stress respective to the Col-0. Col-0, *SiHDA9-OE1, and SiHDA9-OE2* overexpression lines (T3) were grown in soil for three weeks, and **a)** the RT-PCR result showed the *SiHDA9* expression only in both overexpressing transgenic lines but not in Col-0 (wild-type). **b)** The transpirational water loss was measured in four weeks of grown seedlings of both *SiHDA9* overexpression lines and Col-0. The water loss in the detached shoots was calculated as a percentage of the initial fresh weight. Error bars represent the average ± SE five shoots of different plants. **c)** The three weeks soil-grown Col-0 and *SiHDA9* overexpression lines were subjected to 14 days of drought treatment before rewatering. Plants were then allowed to recover for one day. Pictures were captured after 10 and 14 days of drought and after one day recovery. **d)** The RWC (relative water content) was also measured in the watered and drought-treated (10 days) Col-0 and *SiHDA9* overexpressing plants. The result showed a significantly lower RWC in *SiHDA9* overexpressing plants of both the lines in drought conditions respective to the Col-0 plants, whereas, no significant differences were observed between them in watered conditions. **e)** Survival rate of 14 days drought-stressed plants after rewatering for one day showed zero percent recovery in the *SiHDA9* overexpressed plants of both lines, whereas it was 50% in the Col-0. Error bars represent the standard error of three biological replicates. Asterisks represented the student’s t-test: * *P* <0.05 and ** *P* <0.01.

Transpirational water loss is an utmost factor associated with drought tolerance. The transgenic lines were grown in soil to assess the variations in the rate of water loss due to overexpression of SiHDA9. There were no significant observations in the phenotypic parameters (leaf number, leaf length, and leaf width) except rosette diameter (Extended Fig. 12a-d). The fresh weight changes in the detached rosettes of these lines were measured for 200 minutes. The *SiHDA9* overexpression lines showed significantly faster water loss in contrast to the wild-type (Fig. 4b). This suggested the decreased drought tolerance in overexpression lines compared to WT plants. Further, the overexpression lines with Col-0 were subjected to the drought tolerance assay to elucidate the negative role of SiHDA9 in the dehydration stress tolerance. We observed that plants of both the *SiHDA9* overexpression lines were more wilted under drought conditions (10 and 14 days) imposed by withholding water for 14 days (Fig. 4c). Next, the relative water content (RWC) was measured to confirm the phenotypic observations (wilting) in the drought tolerance assay. Both the overexpression lines, *SiHDA9-OE1* (58.3 ± 9.28 %) and *SiHDA9-OE2* (51.9 ± 10.18 %), displayed a significant (*P* < 0.05) lesser RWC than Col-0 (72.5 ± 3.84) after 10 days of drought treatment. There were no significant variances in their well-watered control (Fig. 4d). Next, to test the survival rate of plants, the 14 days drought-treated plants were rehydrated for one day. All the plants of both the transgenic lines did not recover, while 50 ± 10.2 % of plants survived in the case of wild-type (Fig. 4e). These results validated that SiHDA9 plays a negative role in drought-stress tolerance.

### SiHDA9 physically interacts with SiHAT3.1 and SiHDA19

To investigate the mechanistic details of SiHDA9, we decided to identify the associated protein complex because HDA9 acts in the multiprotein complex to regulate different biological processes^12,15,16,19–25^. Initially, the co-expression correlation analysis was performed and determined the co-expressing gene network with *SiHDA9* using the RNA-seq data of PEG treatments at different time points (0, 6, 12, and 24 h) in both the IC4 and IC41 cultivars. The analysis identified 2293 positively co-expressed genes (PEGs) and 285 negatively co-expressed genes (NEGs) with *SiHDA9* in IC4, whereas the 2365 PEGs and 846 NEGs with *SiHDA9* in IC41 cultivar (Supplementary Table 8). The collective expression analysis of *SiHDA9* PEGs and NEGs in the IC41 cultivar showed a linear increasing and decreasing expression pattern from 0 to 24 h of PEG treatments (Fig. 5a,b), respectively. In case of the IC4, there was no linearity found in the expression pattern of both PEGs and NEGs (Extended data Fig. 13a,b). Next, the pathway analysis of PEGs and NEGs of both cultivars were also performed and recognized the more significant enrichment of stress-responsive pathways in the NEGs of IC41 (Extended data Fig. 13c, Supplementary Table 9). These results indicated that SiHDA9 might suppress the expression of stress-responsive genes in IC41. Thus, the STRING tool predicted the interacting partners of SiHDA9 in the PEGs in both cultivars. The investigation showed that the SiHDA9 could physically interact with different proteins, including, SiHAT3.1, SiHD1 (SiHDA19), SiMSI1, SiELF6, SiHAG4, SiHAG1 and SiMRG in IC41 (Fig. 5c), whereas only with SiMRG, SiPKL and SiCHR4 in the case of IC4 cultivar (Extended data Fig. 13d). It was previously reported that HDA9 acts in the form of a repressor complex consisting of AtHDA9, PWR and WRKY53 which promoted aging in Arabidopsis^20^. SiHAT3.1 is a PHD-finger homeodomain protein^40^ that showed the highest co-expression correlation score (0.9908) among all the predicted interacting partners with SiHDA9. So, we speculated that SiHAT3.1 might help to recruit a whole complex on the DNA by its DNA binding motif. To confirm this supposition, the yeast-two hybrid assay (Y2H) was performed to validate the interactions of SiHDA9 with SiHAT3.1. SiHDA19 was also included as a positive control in this study. It was also found as an interacting partner of HDA9 in Arabidopsis^12^ and up-regulated in both the foxtail millet cultivars. Y2H assay showed that SiHDA9 could interact with SiHAT3.1 and SiHDA19 (Fig. 5d). Further, to validate these interactions *in planta*, we performed the Bimolecular fluorescence complementation assay (BiFC). For the BiFC, the SiHDA9 was fused to N-terminal, whereas SiHAT3.1 and SiHDA19 were fused to the C-terminal halves of Yellow Fluorescent Protein (YFP). These constructs were transformed in the *Agrobacterium* and coinfiltrated in tobacco leaves (*Nicotiana benthamiana*). With the desired pair of proteins, the nuclear localization signal (NLS) containing marker protein with Red Fluorescent Protein (RFP) tag was also co-infiltrated to visualize the nucleus. The confocal microscopy of the agro-infiltrated leaves showed YFP signals in the nucleus (Fig. 5e). Thus, the BiFC results confirmed that the direct physical interactions *in planta* between SiHDA9-SiHAT3.1 and SiHDA9-SiHDA19 mainly occur in the nucleus.

**Fig. 5.**
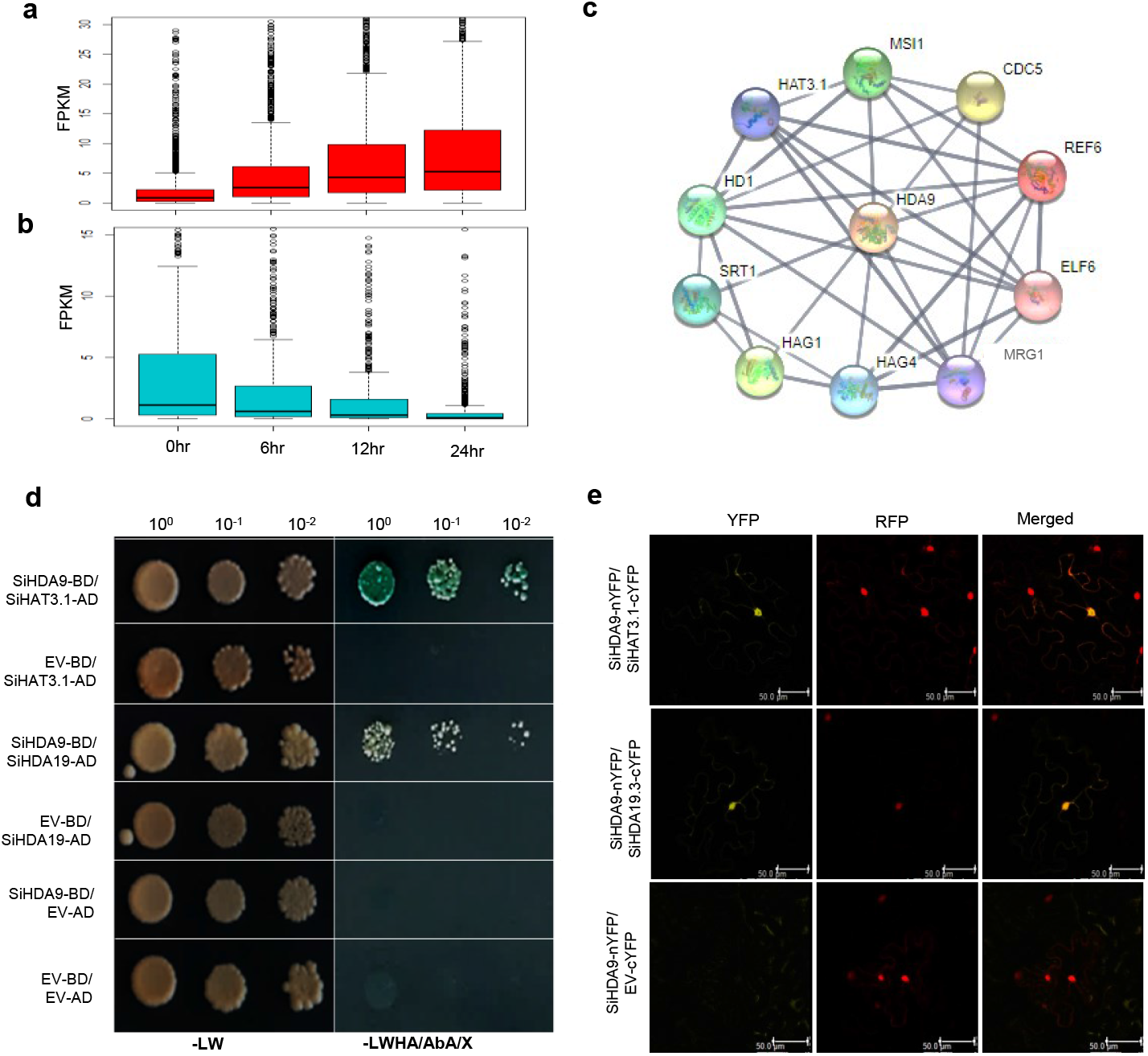
SiHDA9 co-expressed and physically interacts with SiHAT3.1 and SiHDA19. Box plot showing the expression pattern of **a)** Positively co-expressed genes (PEGs) & **b)** Negatively co-expressed genes (NEGs) at 0, 6, 12, and 24hr after PEG treatments in IC41 cultivar. **c)** The string result showed the possible interacting partners of SiHDA9 in the PEGs of the IC41 cultivar of foxtail millet. Proteins are represented by different colored nodes, while the protein-protein association is represented by grey lines or edges at a high confidence level (0.700). **d)** Yeast two-hybrid (Y2H) assays were performed to validate interactions. SiHDA9 fused with the DNA-binding domain (BD) of GAL4, whereas SiHAT3.1 and SiHDA19 with the transcriptional activation domain (AD) of GAL4. The interactions were inspected by cell growth on selective media; double drop-out -LW (- Leu and Trp) and quadruple drop-out -LWHA (- Leu, Trp, His, and Ade) in the plates. EV-AD and EV-BD represent the empty vector (EV) without cloning any gene. AbA represents the antibiotic Aureobasidin A, and X stands for X-α-Gal, which uses to detect α-galactosidase activity for blue color. **e)** Bimolecular fluorescence complementation (BiFC) assays showed protein-protein interactions (SiHDA9-SiHAT3.1 and SiHDA9-SiHDA19) occurred mainly in the nucleus. SiHDA9 fused with the N-terminal while SiHAT3.1 and SiHDA19 with the c-terminal of fragmented YFP protein. The constructs were co-expressed in the tobacco leaves, and YFP signals (yellow color) were scanned on a confocal microscope after 48hrs of Agroinfiltration. IDD14-RFP (red color) used as a nucleus marker. Scale bar, 50 μm.

### SiHDA9-HAT3.1-SiHDA19 complex recruited on the promoter of drought-responsive genes to repress their expression

The findings that SiHDA9 and SiHAT3.1 co-expressed in dehydration conditions and interacted in the nucleus prompted us to identify the target genes. The pathway analysis of our integrative data of ChIP-seq and RNA-seq showed enrichment of several stress-related pathways, including response to water deprivation, which was restricted in the only cUP-gUP category but not in the cDN-gDN of IC4 cultivar. The enrichment of these stress-related pathways was not only restricted in the cUP-gUP category, but it was also more enriched in the cDN-gDN category of IC41 (Fig. 3c). Next, the uniquely and commonly correlated genes between IC4 and IC41 cultivars were identified in our integrative data. The different stress-responsive pathways, including water-deprivation, were more enriched in the unique IC4-cUP-gUP and the unique IC41-cDN-gDN (Supplementary Table 10). Therefore, we assumed that the drought-responsive genes of these categories might be important for drought tolerance. Thus, to identify the SiHDA9 target genes, the promoters of unique stress-responsive genes (including water deprivation encoding genes) of both categories were analysed for the presence of the SiHAT3.1 binding conserved motif (T(A/G)(A/C)ACCA) (Fig. 6a). The motif enrichment analysis identified a total of 42 out of 68 and 21 of 36 genes containing this motif in the unique IC4-cUP-gUP of IC4 and unique IC41-cDN-gDN of IC41 categories, respectively (Supplementary Table 11). These genes include *SiRAB18, SiRAP2.4*, and *SiP5CS2* in unique IC4-cUP-gUP and *SiRD22, SiPIP1;4*, and *SiLHCB2.3* in unique genes of IC41-cDN-gDN. These genes showed higher expression levels (Log2 Fold Change) after 24 h PEG treatment in IC4 compared to the IC41 cultivar (Extended data Fig. 14a,b). Thus, the H3K9ac peak enrichment in these genes was visualized by IGV (Integrative Genome Visualizer) in both cultivars. Interestingly, in the case of all six genes, we detected the depleted H3K9ac peak (lower acetylation level) around the HAT3.1 recognition site (on the promoter region) in IC41-P (24 h PEG) sample in comparison to the control sample (IC41-C) (Fig. 6b,c). In contrast, the peak around the SiHAT3.1 binding motif was enriched in the case of all the genes compared to the IC4-C (control sample). Furthermore, to check the impact of deacetylation on the expression of these genes, we performed the inhibitory assay experiment. In this, seedlings were treated with trichostatin A (TSA), a histone deacetylase (HDAC) inhibitor. The expression of *SiRAB18* and *SiP5CS2* was induced after 24 h of TSA treatment in both cultivars. While the expression of *SiRAP2.4* was reduced than the control, it was higher in IC4 respective to the IC41 cultivar (Extended data Fig. 15). So, these results suggested that histone deacetylases might regulate these genes. Hence, overall results depicted that SiHDA9 with SiHAT3.1 might directly regulate the expression of these genes in the sensitive cultivar (IC41). To confirm these results, the expression of homologs of these genes was tested in the *SiHDA9* overexpression lines in Arabidopsis. Expression analysis by qRT-PCR showed the higher induction of *AtRAB18, AtRAP2.4, AtP5CS2, AtRD22, AtPIP1;4* and *AtLHCB2.3* in the Col-0 after drought treatment. Whereas these genes fail to attain the expression level in both the overexpression lines (*SiHDA9-OE1 and SiHDA9-OE2*) after drought treatment, as found in the Col-0 (Fig. 6e-j). Thus, these results demonstrated that the overexpression of *SiHDA9* might be inhibited the expression induction of drought-responsive *SiRAB18, SiRAP2.4, SiP5CS2 SiRD22, SiPIP1;4* and *SiLHCB2.3* genes in IC41 cultivar and make it sensitive towards drought.

**Fig. 6.**
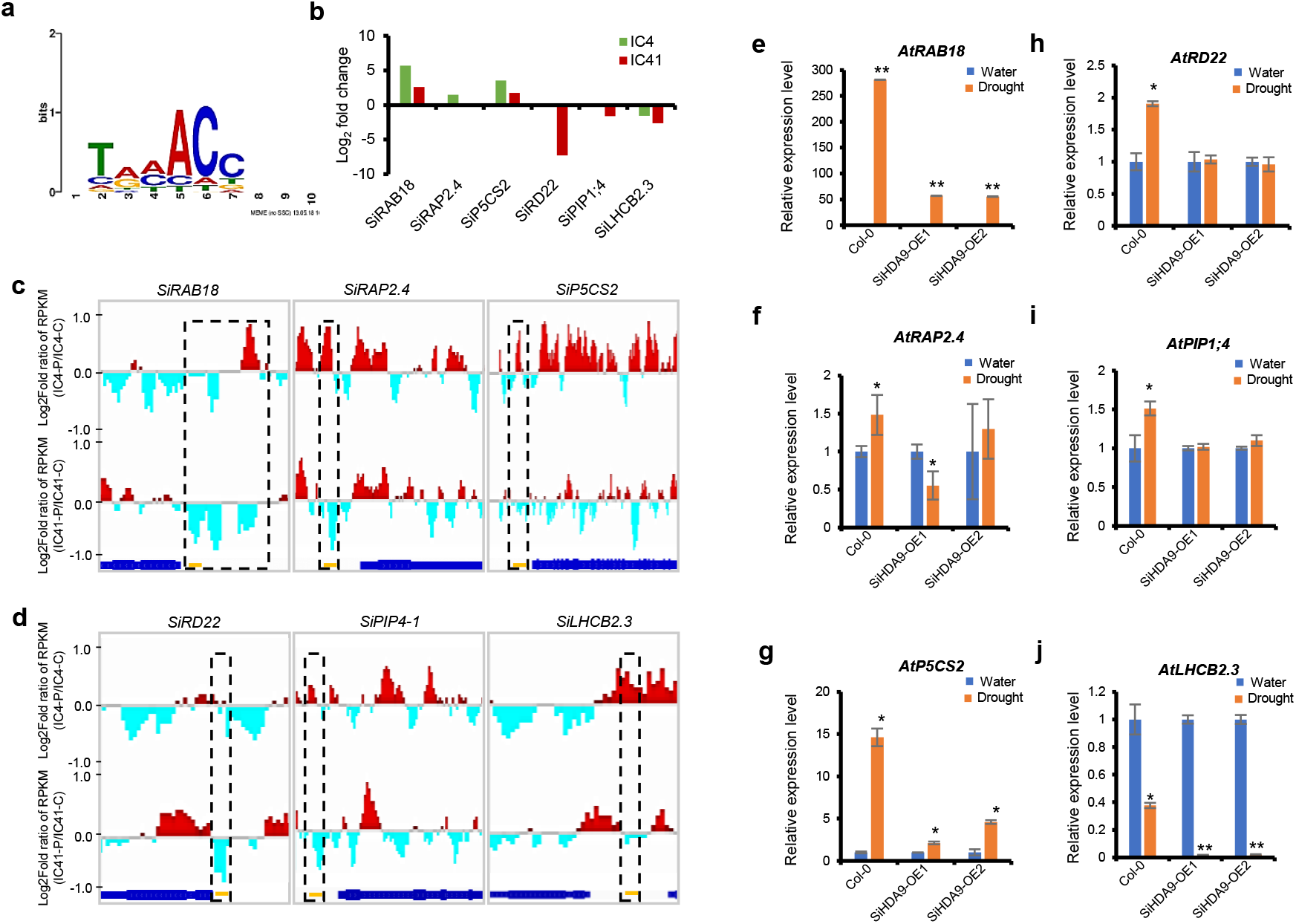
SiHDA9-SiHDA19-HAT3.1 repressor complex suppresses the expression of drought-responsive genes by decreasing the H3K9ac levels on their promoter after 24hr PEG treatment in the sensitive cultivar of foxtail millet. **a)** Conserved binding motif of HAT3.1 present in the water-deprivation encoding genes including *SiRAB18, SiRAP2.4* and *SiP5CS2* in unique IC4-cUP-gUP *and SiRD22, SiPIP4-1 and SiLHCB2.3* in unique IC41-cDN-gDN. **b)** These genes were significantly differentially expressed (Log2Fold Change) after PEG-treatment present in the RNA-seq. data. IGV screenshots of these representative genes showing the distribution of H3K9ac level in the term of Log2 fold of RPKM (+1 to −1) at the **c)** unique IC4-cUP-gUP genes; *SiRAB18, SiRAP2.4, SiP5CS2*, and **d)** IC41-cDN-gDN; *SiRD22, SiPIP4-1*, and *SiLHCB2.3* genes. Orange-colored dashes represent the HAT3.1 binding motif on the promoter of the genes. The black colored (dashes) box represents the differential peaks around the HAT3.1 binding motif. RPKM stands for Reads Per Kilobase of the transcript, per Million mapped reads. Expression pattern of **e)** *AtRAB18*, **f)** *AtRAP2.4*, **g)** *AtP5CS2* **h)** *SiRD22*, **i)** *SiPIP4-1* and ***j*)** *SiLHCB2.3* in *SiHDA9* overexpression lines in Arabidopsis. Error bars represent the standard error between biological replicates. Asterisks represent the student’s t-test: * *P* <0.05 and** *P* <0.01.

## Discussion

It has been known that the H3K9ac level onto the promoter of drought-responsive genes is enriched by GCN5 under drought-stress conditions to confer drought tolerance^5^. But, which histone deacetylase (HDAC) antagonized this action is unclear. It has also been studied that HDA9 negatively regulates the mechanism of salt and drought stress tolerance in Arabidopsis^12^. It repressed the expression of stress-responsive genes by regulating the H3K9ac level on their promoter. Various reports highlight that HDA9 acts in the multiprotein complexes and repress their target genes’ expression to regulate different biological pathways^12,15,16,19–25^. However, the precise molecular mechanism of HDA9 action in regulating drought-responsive genes in plants is not yet fully understood. The current study supports fillings these significant knowledge gaps and provides a better understanding of the drought regulatory mechanisms vital for plant growth and their adaptation to adverse conditions. In this study, we described the SiHDA9 regulatory mechanism that suppressed expression of stress-responsive genes such as *SiRAB18, SiRAP2.4, SiP5CS2, SiRD22, SiPIP1;4* and *SiLHCB2.3* through deacetylation of H3K9ac marks to negatively regulates the drought tolerance in foxtail millet.

The present study identified a higher enrichment of H3K9ac around the TSS in both the cultivars (Extended data Fig. 5) in agreement with previous studies where the enrichment of H3K9ac highly occurred around the TSS^41^. We also observed a higher enrichment of H3K9ac level on the promoter of the drought-responsive genes in the dehydration tolerant cultivar (IC4) after PEG treatment. In contrast, its level decreases after PEG treatment in the sensitive cultivar (IC41) compared to their respective control (Fig. 3d,g and Fig. 6c,d). Consistent with our results, it has been known that the H3K9ac level increases onto the promoter of stress-responsive genes during drought conditions, leading to the gene activation to confer stress tolerance^5,6^. Stress-responsive genes, such as *RAB18, RAP2.4*, and *P5CS2*, were reported to be induced under different stresses, including drought, and their expression was activated through the augmentation of H3K9ac on their promoter^6,9^. We also observed relatively higher expression levels of these drought-responsive genes after PEG treatment in the IC4 compared to the IC41 (Fig. 6b), depicting the role of these genes in conferring dehydration tolerance in the IC4 cultivar.

The study also revealed that the chromatin organization pathway was enriched in the sensitive cultivar compared to the tolerant cultivar (Fig. 2a,b). The genes involved in this pathway include *SiHDA9* and *SiHAT3.1*. SiHDA9 is known to suppress gene expression^12,15,16,19–25^. SiHDA9 is more expressed after 24 h dehydration stress in the dehydration-sensitive cultivar (IC41). The integrative analysis of RNA-seq and ChIP-seq and their pathway analysis also showed that the different stress-related pathways, including response to water deprivation, were significantly more enriched in the cDN-gDN than the cUP-gUP category of IC41. In contrast, these pathways were uniquely enriched in the cUP-gUP category of IC4 (Fig. 3c). These observations suggested that SiHDA9 might be causing the suppression of different stress-responsive pathways encoding genes to decrease the stress responses in the IC41 cultivar. Consistent with these results, transgenic overexpression lines of *SiHDA9* in Arabidopsis showed severe dehydration or drought sensitivity and increased water loss in contrast to the wild-type plants (Fig. 4b-d, Extended data Fig. 11b). Thus, these results suggest that HDA9 is a negative regulator of drought tolerance in foxtail millet, as reported earlier in the case of Arabidopsis^12,17^.

Moreover, results also showed that *SiHDA9* co-expressed and could interact with different epigenetic modifiers encoding genes, including *SiHAT3.1, SiHDA19, SiSRT1, SiHAG1, SiHAG4, SiMRG, SiELF6, SiREF6* and *SiCDC5* (Fig. 5c) in the sensitive cultivar of foxtail millet. We also validated that SiHDA9 physically interacted with the SiHAT3.1 and SiHDA19 (Fig. 5d,e). SiHAT3.1 is a PHD-finger homeodomain box protein and has the specificity to recognize a unique DNA sequence motif (T(A/G)(A/C)ACCA) in Arabidopsis^40,42^. So, it might help recruit repressor complex (SiHDA9-SiHAT3.1-SiHDA19) onto the target genes. In agreement with the previous study in Arabidopsis^12^, we also found interactions between SiHDA9 and SiHDA19. It has been reported that HDA19 also negatively regulates stress tolerance^43^. Therefore, it confirms that SiHDA9 acts as a multiprotein complex which was in agreement with several previous studies^12,15,16,19–25^. The interaction network also showed that the SiHDA9 could interact with the histone acetyltransferase, HAG1 and HAG4 (Fig. 6b). However, SiHDA9 is negative while HAG1 (GCN5) is a positive regulator of drought tolerance^12,5^. There might be a possibility, SiHDA9 physically interacted with HAG1/HAG4 to inhibit their activity (important for gene activation) by deacetylation. Similar to this hypothesis, a previous report showed that HDA9 could also deacetylate to non-histone protein WRKY53 to inhibit its activity, required for the gene activation in Arabidopsis^17^.

Furthermore, the integrative analysis of RNA-seq and ChIP-seq showed that the cUP-gUP category of IC4 and cDN-gDN category of IC41 included different stress-responsive pathways encoding genes (Fig. 3c). Further, the SiHAT3.1 binding motif searched in the stress-responsive genes including water-deprivation pathway encoding genes in the categories unique IC4-cUP-gUP and unique IC41-cDN-gDN which possessed the HAT3.1 binding motif in their promoter (Supplementary Table 10, 11). The previous studies reported that HDA9 preferentially binds on the promoter of active genes to repress their expression^20,23^. In the present study, we found important genes such as *SiRAB18, SiRAP2.4* and *SiP5CS2* in the unique IC4-cUP-gUP and *SiRD22, SiPIP1;4* and *SiLHCB2.3* in the unique IC41-cDN-gDN categories encoding to the water deprivation pathway. These genes show a relatively lower expression level in the IC41 (Fig. 6b, Extended data Fig. 14). The *RAB18, RAP2.4, P5CS2* and *RD22* genes are drought marker genes that provide drought tolerance^44–47^. It is also known that H3K9ac increases on the promoter of RAB18 and RAP2.4 to induce their expression upon stresses encounter ^9,6^. RAB18 is a dehydrin protein and highly induced under different stress conditions, including drought and provides tolerance^48,49^. RAP2.4 is a transcription factor that regulates cuticular wax biosynthesis and is vital in conferring drought tolerance^45^. P5CS2 is a proline (osmoprotectant) biosynthesis encoding gene, important for drought tolerance^39^. RD22 is up-regulated in drought conditions and plays a crucial role in the stomatal closure^50^. PIP1;4 is a water channel (aquaporin) important role in drought tolerance^51^. It is also known that the down-regulation or disruption of the LHCB2.3 protein decreases drought tolerance in Arabidopsis^52^. We observed a lower level of acetylation in the promoter region of these genes after PEG treatment in the IC41 cultivar (Fig. 3g and Fig. 6c,d). In addition, results also showed that the *SiHDA9* overexpression lines in Arabidopsis also failed to induce the expression level of *RAB18, RAP2.4, P5CS2, RD22, PIP1;4*, and *LHCB2.3* genes after drought treatment as found in wild-type Col-0 (Fig. 6e-g). Thus, results depicting that drought-responsive genes, including *SiRAB18, SiRAP2.4, SiP5CS2 SiRD22, SiPIP1;4* and *SiLHCB2.3* were important for the drought tolerance in the IC4, whereas a lower level of their reduced expression might be responsible for the dehydration sensitivity in the IC41 cultivar of foxtail millet (Fig. 7). Overall, the study reveals the epigenetic mechanism of dehydration stress tolerance in foxtail millet, emphasising the functional role of SiHDA9. This knowledge could further be used to impart durable tolerance to dehydration stress in cultivated varieties using gene manipulation and/or genome editing approaches.

**Fig. 7.**
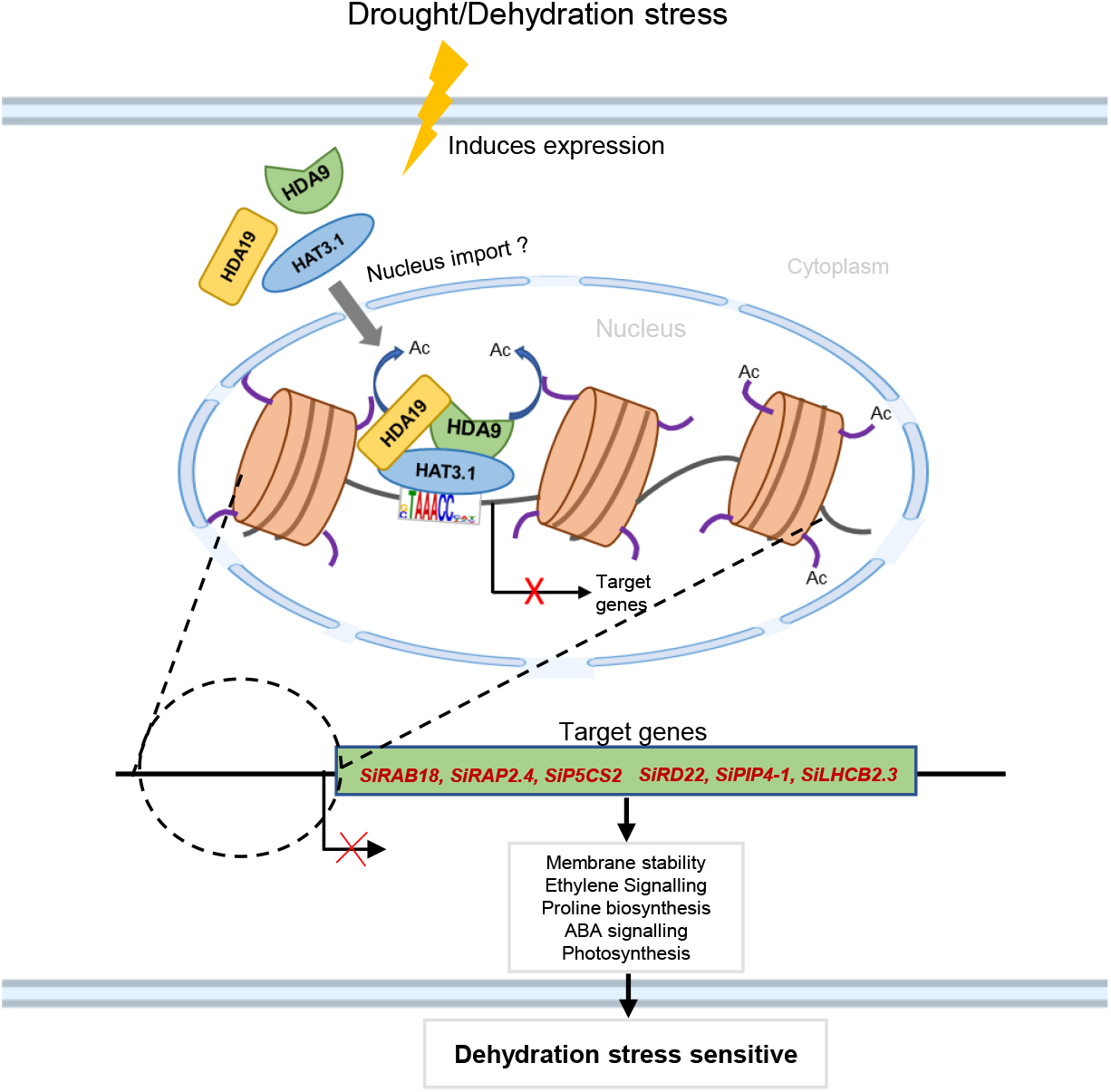
A working model for the biological functions and molecular mechanisms of SiHDA9 in foxtail millet. SiHDA9, SiHAT3.1, and SiHDA19 were up-regulated in response to the dehydration stress and formed a multiprotein repressor complex (SiHDA9-SiHAT3.1-SiHDA19) through physical interaction with each other in the nucleus. This complex recruited on the promoter of drought-responsive targets genes such as *SiRAB18, SiRAP2.4, SiP5CS2, SiRD22, SiPIP4-1, and SiLHCB2.3* by recognizing the SiHAT3.1 binding motif (T(A/G)(A/C)ACCA) and remove the H3K9ac marks to suppress their expression. This might be decreasing membrane stability, ethylene signaling, proline biosynthesis, ABA signaling, and photosynthesis resulted more dehydration sensitivity in the IC41 cultivar of foxtail millet. Further, these genes may induce the de-repression of their downstream target genes to increase the dehydration sensitivity.

## Materials and Methods

### Plant materials

The seeds of salt and dehydration tolerant ‘IC-403579’ (IC4) and sensitive ‘IC-480117’ (IC41) foxtail millet cultivars were used in this study^32,38^. The seeds were sown in a plant growth chamber (PGC-6L; Percival Scientific Inc., USA) with the following conditions; 28±1°C day/23±1 °C night/70±5% relative humidity with a photoperiod of 14 h and a photosynthetic photon flux density of 500 μmol m^−2^ s^−1^ used as in the previous studies^32,38^. For the dehydration stress, the twenty-one days old seedlings were treated with 20% polyethylene glycol 600 (PEG-6000) ^32,38^. The dehydration-stressed samples were collected after 24 h PEG treatments. The untreated samples were also collected as controls (0 h). All harvested samples were frozen immediately in liquid nitrogen and stored at −80 °C until RNA and nuclei were isolated.

### Expression analysis by RNA-Sequencing and qRT-PCR

The total RNA was extracted from PEG-treated (24 h) and untreated control (0 h) samples using Spectrum plant total RNA Kit (Sigma Aldrich, St. Louis, MO, USA) according to the manufacturer’s instructions. The quality of total RNA was checked on the 1% agarose gel. The 10μg of total RNA was treated with DNase (Ambion, Foster City, California, USA) for the removal of DNA contamination. The 5μg total RNA (DNase-treated) was outsourced for the RNA-Seq on Hi-Seq 2000 genome analyzer (Illumina) sequencing platform. The paired-end RNA-Seq was performed for all the samples. Three biological replicates were sequenced for each 0 and 24 h PEG treatment. For qRT-PCR (Quantitative Real-Time PCR), the first-strand cDNA was synthesized using 2μg of DNase-treated total RNA as a template and superscript II from the reverse transcription kit (Invitrogen, Carlsbad CA, USA) follow to the manufacturer’s instructions. The forward and reverse primers were designed by the Primer Express tool. The qRT-PCR amplification was carried out using the ABI Prism 7500 Fast sequence detection system (Applied Biosystems, Foster City, CA, USA) with Fast SYBR™ Green PCR Master Mix (Applied Biosystems™, Foster City, CA, USA) according to the manufacturer’s instructions. The fold change was calculated using the 2^−ΔΔCt^ method, and the *SiActin2* gene was used as an internal control for normalization ^29^. Three biological and two technical replicates were used in this experiment.

### RNA-Sequencing analysis

All the raw reads from sequencing data of 0 h (control) and 24 h samples were quality filtered using the FASTX toolkit (http://hannonlab.cshl.edu/fastx_toolkit/). Then, the quality-filtered reads were mapped on the *S. italica* reference genome^28^ using STAR (version 2.5.4) RNA-seq aligner^53^ with default parameters^53^. The FPKM (Fragments Per Kilobase of transcript per Million mapped reads) values were determined by exclusively mapped reads onto the genome for all the samples using cufflinks^54^. The differential expressed genes (DEGs) were identified by the pair-wise comparison between 0 h and 24 h PEG-treated samples using cuffdiff^54^. The significant genes (≥ ±1.0 Log2 Fold change, FDR<0.05) were shown by the volcano plot created by R-package. The Venny 2.1 online tool (https://bioinfogp.cnb.csic.es/tools/venny/) was used to create a Venn diagram to identify common and unique DEGs between IC4 and IC41 cultivars. The pathway enrichment in the DEGs was assessed by AgriGO online tool (http://systemsbiology.cau.edu.cn/agriGOv2). To determine the clusters (Temporal co-regulation of transcripts) enrichment between IC4 and IC41 cultivars, we calculated the Log2 ratio (IC41/IC4) and used it in the Short Time-Series Expression Miner (STEM) tool^55^. To detect the correct and significant (Bonferroni-corrected *P* <0.05) gene cluster, we applied a minimum 0.7 correlation coefficient and permitted up to 50 permutations during the analysis. The full parameters of the STEM model are given in Supplementary Table 6. The heatmap for the gene expression was generated by the MeV tool^56^. In this study, the boxplots were generated by the GraphPad prism tool.

### ChIP-Sequencing

We determined the differences in the H3K9ac pattern between 0 h and 24 h PEG-treated samples by Chromatin immunoprecipitation (ChIP) assay. The ChIP assay was performed according to the previously described protocol^57^ with some modifications. In brief, the 0 h and 24 h PEG-treated samples (4 g) of both IC4 and IC41 cultivars were crosslinked, followed by crushing to the fine powder using liquid nitrogen. Further, this powder was resuspended in the nuclei isolation buffer, homogenized and filtered through four layers of cheesecloth. The filtered slurry was centrifuged at 11,000g for 20 min at 4°C. Then the nuclei pellet was resuspended in nuclei lysis buffer and further subjected to chromatin sonication. After sonication, these samples were centrifuged at 13,800g for 10 min at 4°C, and supernatant (sonicated chromatin) was collected and used for the ChIP assay. The collected supernatant was pre-cleared by mixing 40 μl Protein A agarose beads (Invitrogen, Carlsbad CA, USA) and further incubated for 1 h at 4°C with gentle agitation. Then, the slurry was centrifuged at 16000 g for 2 minutes at 4°C and collected the supernatant. This supernatant was divided into ChIP, input control, and no-antibody control (No-Ab, negative control). The ChIP samples were immunoprecipitated by anti-H3K9Ac antibody (Cat. No. 06-942, Millipore) using 60 μl protein A agarose beads followed by washing with low and high salt, LiCl and TE buffer. Then, the ChIP-DNA was eluted by freshly prepared elution buffer (1% SDS and 0.1M NaHCO_3_)^18^ followed by reverse crosslinking overnight at 65°C after adding 20 μl of 5M NaCl. Next, the proteinase K treatment was performed at 55°C for 1 h followed by DNA purification. Then, ChIP and input DNA samples of both cultivars were outsourced for sequencing.

### ChIP-Sequencing analysis

The sequenced reads of 0 h (control) and 24 h samples were quality filtered using Trim Galore (https://www.bioinformatics.babraham.ac.uk/projects/trim_galore/). Then, the quality-filtered reads were mapped on the *S. italica* reference genome^28^ using Burrows-Wheeler Aligner (BWA) with default parameters^58^. Next, the peak calling was performed in the ChIP samples against sequencing reads from the input sample by MACS2 (version 2.1.4)^59^ to identify the enriched H3K9ac peaks (FDR value <0.05). Further, these enriched peaks were annotated for the genomic regions by HOMER (http://homer.ucsd.edu/homer/ngs/annotation.html). Next, we investigate the correlation between ChIP-seq and RNA-seq to identify the DEGs due to the modulations in the H3K9ac mark. For this, the bam file of ChIP-seq and expression data of RNA-seq were used in Epigenomix software using Bayes MixModel (Z-score) with default parameters^60^ to get the positive (cUP-gUP and cDN-gDN) and negative (cUP-gDN and cDN-gUP) correlated genes within ±2kb of the TSS with default parameters. The pathway enrichment analysis in the cUP-gUP and cDN-gDN categories of both cultivars was performed by AgriGO v2. Whereas scatter plots were generated by R-package. Motif scanning was performed by FIMO (Find Individual Motif Occurrences) online tool (https://meme-suite.org/meme/tools/fimo).

### Co-expression correlation analysis

To determine the co-expression gene network of *SiHDA9*, we have used the RNA-seq data of 0, 6, 12^31^ and 24 h PEG-treated samples of IC4 and IC41. The Log2 transformed FPKM (Fragments Per Kilobase of transcript per Million mapped reads) values of all the datasets have been intended by cufflink. These values were further used as input in Cytoscape version 2.8.1 (http://apps.cytoscape.org/apps/expressioncorrelation) to identify the co-expressed genes with *SiHDA9*. Based on Person’s correlation coefficient (r), the positive; PEGs (r ≥ 0.95) and negative; NEGs (r ≤ −0.95) co-expressed genes with *SiHDA9* were identified at different time points of PEG treatments. The cumulative expression pattern of these genes is shown by Boxplots, which were constructed by R-package. Furthermore, the pathways analysis in the *SiHDA9* PEGs and NEGs of both cultivars were performed by AgriGO v2, whereas protein-protein interaction network analysis in the PEGs with *SiHDA9* in both cultivars was performed by String online tool (https://string-db.org/) at a high confidence level (0.700).

### Yeast two-hybrid assay

The method of yeast two-hybrid (Y2H) assays used in this study has been previously described^61^. The full-length coding sequence (CDS) of the *SiHDA9, SiHAT3.1* and *SiHDA19* has been amplified by the primers mentioned in Supplementary Table 12 and cloned in the yeast expression vectors. *SiHDA9* was cloned in the pGBKT7-BD (bait vector), whereas *SiHAT3.1* and *SiHDA19* were cloned in the pGADT7-AD vector (pray vector). The constructs were transformed in the Y2H gold yeast strain by using Yeastmaker Yeast Transformation System (Clontech). For the Y2H assay, the yeast cells containing pairs of BD and AD-derived constructs were grown on a medium with DDO (-LW; Leu and Trp) and QDO (-LWHA; Leu, Trp, His, and Ade). QDO was also supplemented with antibiotics Aureobasidin A (AbA) and X-α-Gal (for blue color development).

### Bimolecular fluorescence complementation (BiFC) assay

For the BiFC assay, the coding sequence of *SiHDA9, SiHAT3.1*, and *SiHDA19* has been amplified by the primers mentioned in Supplementary Table 12. The Gateway cloning system was used for the cloning of the genes into the vectors pSITE-N (*SiHDA9*) and pSITE-C (*SiHAT3.1* and *SiHDA19*) to express protein fusions with N- or C-terminal domain of the yellow fluorescent protein (YFP), respectively. The *Nicotiana benthamiana* leaves were co-infiltrated with *Agrobacterium tumefaciens* (strain GV3101) containing pSITE-N and pSITE-C derived constructs. With both the constructs, additionally, the NLS (nuclear localization signal) fused with RFP (red fluorescent protein) tag-containing construct was also co-infiltrated to spot the nucleus. After 48 h of infiltration, the images were captured on the Leica TCS SP5 confocal spectral microscopy imaging system.

### Dehydration stress tolerance assay in yeast

For the dehydration tolerance assay in yeast, first, we cloned the *SiHDA9* in the yeast expression vector pYES2. Then, pYES2-SiHDA9 and empty pYES2 vector (EV) were transformed individually into Saccharomyces cerevisiae strain W303 using Yeastmaker Yeast Transformation System (Clontech). The transformed cells were screened on SD/-ura medium with 2% (w/v) dextrose at 30°C for 72 h and used in the abiotic stress tolerance assay described previously^29^. For the dehydration stresses, yeast cells harboring pYES2-SiHDA9 and pYES2-EV were grown in SD/-ura medium supplemented with 30% PEG-6000 (polyethylene glycol-6000) and incubated at 30°C for 36 h. After that, the stressed and unstressed (for control) cultures were spotted (with serial dilutions) on the basal SD/-ura medium plates, supplemented with 2% (w/v) dextrose and incubated at 30°C for 72 h.

### Generation and analysis of Arabidopsis transgenic lines

For the overexpression, we cloned a full-length coding sequence of *SiHDA9* in the pGWB420 vector (https://www.addgene.org/74814/). The prepared construct (CaMV35S: *SiHDA9*) was introduced into the Col-0 using the floral dip transformation method in Arabidopsis^62^. The transgenic lines were screened by antibiotics selection followed by *nptII* gene-specific PCR. The expression analysis by qRT-PCR was performed in the T2 and T3 generation. The T3 lines were also analyzed for the water loss, drought tolerance assay and relative water content (RWC) estimation, and stress recovery.

### Dehydration/drought stress tolerance assays in Arabidopsis

To evaluate the impact of dehydration stress in *SiHDA9* overexpression lines (*SiHDA9-1* and *SiHDA9-2*) and Col-0, the stress treatment was given by the mannitol as described previously^49^. The seedlings from *SiHDA9* overexpression lines and Col-0 were cultured in ½ Murashige and Skoog (MS) medium for one week and then shifted into plates containing ½ MS (control) or ½ MS augmented with 150mM and 300mM concentrations of mannitol and permitted to grow vertically for 14 days. The seedling’s growth (biomass) was measured and photographed. The experiment was performed in triplicates.

For the drought treatment, seeds of *SiHDA9* overexpression lines and Col-0 were sown in the pots and provided normal watering conditions. After 21 days, the water supply was suspended until 14 days to create the drought stress, and then, drought-treated plants were rewatered, and recovery was estimated after 24 h^63^. Drought stress tolerance assays were performed in duplicate, but only one representative image is shown. Four biological replicates were used for the drought experiment. The relative water content (RWC) was also measured in the 10 days of drought-treated plants. For this, shoots of *SiHDA9* overexpression lines and Col-0 camta1-2 and camta1-3 were detached and weighed instantly to record fresh weight (FW); then, shoots were dipped in distilled water for 4 h. After that shoot were blotted and weighed to record the turgid weight (TW). The dry weight (DW) was also calculated after drying the turgid shoots at 80°C for 24 h. The RWC was determined by this formula; RWC (%) = (FW-DW) / (TW-DW) X 100^38^. For the water loss estimation, the shoots of 4-week-old plants (well-watered) were detached from the root system and weighed instantly for the fresh weight (initial weight). Then, the shoots were placed in Petri dishes on a laboratory bench and followed the weighing at different intervals; 20, 40, 60, 80, 100, 120, 160, and 200 minutes^63^. The lost proportion (percentage) at different time intervals was calculated based on the fresh weight of the rosettes of individual plants.

### Statistical Analysis

The mean value of data from the independent experiment is shown in the figures. The errors bars shown in the graphs are the standard error. The statistical significance was calculated using the unpaired student’s t-test via excel and GraphPad software. The significance level was shown by asterisk after calculating the *p-value (*P* < 0.05; \*\**P* < 0.01; \*\*\**P* < 0.001) between control and treated samples.

## Data availability

The data presented in this study are deposited in the National Center for Biotechnology Information (NCBI) Sequence Read Archive (SRA) database under BioProject PRJNA744331 and PRJNA734765 for RNA-seq and PRJNA744019 for ChIP-seq.

## Authors contributions

M.P. and V.K. conceived the idea and planned the work. V.K. performed maximum of the experiments including RNA-seq, drought stress related experiments in foxtail millet and Arabidopsis, Co-expression correlation analysis, Y2H, BiFC assay, RT-PCR etc. V.K. with the assistance of N.S. performed ChIP-assay. V.K. and B.S. performed bioinformatic data analysis of ChIP-seq and RNA-seq and dehydration stress assay in Arabidopsis. B.S. and S.V.S. helped in data analysis and its interpretation. V.K. wrote the manuscript. M.M. and M.P. helped in the data interpretation and constructing the manuscript.

## Acknowledgements

Authors’ work in this area is supported by J.C. Bose National Fellowship Grant of Department of Science and Technology, Government of India (File No.: JCB/2018/000001) and Core Grant of National Institute of Plant Genome Research, New Delhi, India. V.K. acknowledges the Science and Engineering Research Board, Government of India, for National Post-Doctoral Fellowship Grant (File no. PDF/2017/000892). The authors thank DBT-eLibrary Consortium (DeLCON) for providing access to e-resources. We also thank Mr. Anand Dangi for his assistance with experiments.

## References

1. Han, S. K. & Wagner, D. Role of chromatin in water stress responses in plants. J. Exp. Bot. 65, 2785–2799 (2014).

2. Kumar, V., Thakur, J. K. & Prasad, M. Histone acetylation dynamics regulating plant development and stress responses. Cell. Mol. Life Sci. 78, 4467–4486 (2021).

3. Leng, G. & Hall, J. Crop yield sensitivity of global major agricultural countries to droughts and the projected changes in the future. Sci. Total Environ. 654, 811–821 (2019).

4. Kouzarides, T. Chromatin Modifications and Their Function. Cell 128, 693–705 (2007).

5. Li, S. et al. The AREB1 transcription factor influences histone acetylation to regulate drought responses and tolerance in populus trichocarpa. Plant Cell 31, 663–686 (2019).

6. Kim, J. M. et al. Alterations of lysine modifications on the histone H3 N-tail under drought stress conditions in Arabidopsis thaliana. Plant Cell Physiol. 49, 1580–1588 (2008).

7. Kim, J. M. et al. Transition of chromatin status during the process of recovery from drought stress in arabidopsis thaliana. Plant Cell Physiol. 53, 847–856 (2012).

8. Zhang, J. B. et al. A histone deacetylase, GhHDT4D, is positively involved in cotton response to drought stress. Plant Mol. Biol. 1, 1–13 (2020).

9. Kaldis, A., Tsementzi, D., Tanriverdi, O. & Vlachonasios, K. E. Arabidopsis thaliana transcriptional co-activators ADA2b and SGF29a are implicated in salt stress responses. Planta 233, 749–762 (2011).

10. Widiez, T. et al. The chromatin landscape of the moss Physcomitrella patens and its dynamics during development and drought stress. Plant J. 79, 67–81 (2014).

11. Chen, L. T. & Wu, K. Role of histone deacetylases HDA6 and HDA19 in ABA and abiotic stress response. Plant Signal. Behav. 5, 1318–1320 (2010).

12. Zheng, Y. et al. Histone deacetylase HDA9 negatively regulates salt and drought stress responsiveness in Arabidopsis. J. Exp. Bot. 67, 1703–1713 (2016).

13. Song, J., Henry, H. A. L. & Tian, L. Brachypodium histone deacetylase BdHD1 positively regulates ABA and drought stress responses. Plant Sci. 283, 355–365 (2019).

14. Kang, M. J., Jin, H. S., Noh, Y. S. & Noh, B. Repression of flowering under a noninductive photoperiod by the HDA9-AGL19-FT module in Arabidopsis. New Phytol. 206, 281–294 (2015).

15. Khan, I. U. et al. PWR/HDA9/ABI4 Complex Epigenetically Regulates ABA Dependent Drought Stress Tolerance in Arabidopsis. Front. Plant Sci. 11, 623 (2020).

16. Baek, D. et al. Histone Deacetylase HDA9 With ABI4 Contributes to Abscisic Acid Homeostasis in Drought Stress Response. Front. Plant Sci. 11, 143 (2020).

17. Zheng, Y. et al. Histone deacetylase HDA9 and transcription factor WRKY53 are mutual antagonists in regulation of plant stress response. Mol. Plant 13, 598–611 (2020).

18. Kumar, V. et al. Role of GhHDA5 in H3K9 deacetylation and fiber initiation in Gossypium hirsutum. Plant J. 95, 1069–1083 (2018).

19. Tasset, C. et al. POWERDRESS-mediated histone deacetylation is essential for thermomorphogenesis in Arabidopsis thaliana. PLoS Genet. 14, 1–21 (2018).

20. Chen, X. et al. POWERDRESS interacts with HISTONE DEACETYLASE 9 to promote aging in Arabidopsis. Elife 5, e17214 (2016).

21. Lee, K., Mas, P. & Seo, P. J. The EC-HDA9 complex rhythmically regulates histone acetylation at the TOC1 promoter in Arabidopsis. Commun. Biol. 2, 143 (2019).

22. Yang, C. et al. HY5-HDA9 Module Transcriptionally Regulates Plant Autophagy in Response to Light-to-Dark Conversion and Nitrogen Starvation. Mol. Plant 13, 515–531 (2020).

23. Yang, L. et al. HOS15 and HDA9 negatively regulate immunity through histone deacetylation of intracellular immune receptor NLR genes in Arabidopsis. New Phytol. 226, 507–522 (2020).

24. Park, H. J. et al. Hos15 interacts with the histone deacetylase hda9 and the evening complex to epigenetically regulate the floral activator gigantea. Plant Cell 31, 37–51 (2019).

25. Kim, Y. J. et al. POWERDRESS and HDA9 interact and promote histone H3 deacetylation at specific genomic sites in *Arabidopsis*. Proc. Natl. Acad. Sci. 113, 14858–14863 (2016).

26. Bandyopadhyay, T., Muthamilarasan, M. & Prasad, M. Millets for next generation climate-smart agriculture. Front. Plant Sci. 8, 1266 (2017).

27. Lata, C., Bhutty, S., Bahadur, R. P., Majee, M. & Prasad, M. Association of an SNP in a novel DREB2-like gene SiDREB2 with stress tolerance in foxtail millet [Setaria italica (L.)]. J. Exp. Bot. 62, 3387–3401 (2011).

28. Bennetzen, J. L. et al. Reference genome sequence of the model plant Setaria. Nat. Biotechnol. 30, 555–561 (2012).

29. Singh, R. K. et al. Genome-wide analysis of heat shock proteins in C 4 model, foxtail millet identifies potential candidates for crop improvement under abiotic stress. Sci. Rep. 6, 1–14 (2016).

30. Li, C., Yue, J., Wu, X., Xu, C. & Yu, J. An ABA-responsive DRE-binding protein gene from Setaria italica, SiARDP, the target gene of SiAREB, plays a critical role under drought stress. J. Exp. Bot. 65, 5415–5427 (2014).

31. Muthamilarasan, M. et al. Comparative Transcriptome Profiling of Two Contrasting Foxtail Millet Cultivars Provides Insights into Molecular Mechanisms Underlying Dehydration Stress Response. J. Plant Growth Regul. 15, 1–9 (2022).

32. Lata, C., Bhutty, S., Bahadur, R. P., Majee, M. & Prasad, M. Association of an SNP in a novel DREB2-like gene SiDREB2 with stress tolerance in foxtail millet [Setaria italica (L.)]. J. Exp. Bot. 62, 3387–3401 (2011).

33. WeiWei, L. et al. Overexpression of the autophagy-related gene SiATG8a from foxtail millet (Setaria italica L.) confers tolerance to both nitrogen starvation and drought stress in Arabidopsis. Biochem. Biophys. Res. Commun. 468, 800–806 (2015).

34. Singh, R. K., Muthamilarasan, M. & Prasad, M. SiHSFA2e regulated expression of SisHSP21.9 maintains chloroplast proteome integrity under high temperature stress. Cell. Mol. Life Sci. 79, 1–16 (2022).

35. Pandey, G., Yadav, C. B., Sahu, P. P., Muthamilarasan, M. & Prasad, M. Salinity induced differential methylation patterns in contrasting cultivars of foxtail millet (Setaria italica L.). Plant Cell Rep. 36, 759–772 (2017).

36. Yadav, C. B., Muthamilarasan, M., Dangi, A., Shweta, S. & Prasad, M. Comprehensive analysis of SET domain gene family in foxtail millet identifies the putative role of SiSET14 in abiotic stress tolerance. Sci. Rep. 6, 1–13 (2016).

37. Lata, C., Sahu, P. P. & Prasad, M. Comparative transcriptome analysis of differentially expressed genes in foxtail millet (Setaria italica L.) during dehydration stress. Biochem. Biophys. Res. Commun. 393, 720–727 (2010).

38. Lata, C., Jha, S., Dixit, V., Sreenivasulu, N. & Prasad, M. Differential antioxidative responses to dehydration-induced oxidative stress in core set of foxtail millet cultivars [Setaria italica (L.)]. Protoplasma 248, 817–828 (2011).

39. Rymen, B. et al. Histone acetylation orchestrates wound-induced transcriptional activation and cellular reprogramming in Arabidopsis. Commun. Biol. 2, 1–15 (2019).

40. Viola, I. L. & Gonzalez, D. H. Interaction of the PHD-finger homeodomain protein HAT3.1 from Arabidopsis thaliana with DNA. Specific DNA binding by a homeodomain with histidine at position 51. Biochemistry 46, 7416–7425 (2007).

41. Wang, L. et al. Ethylene induces combinatorial effects of histone H3 acetylation in gene expression in Arabidopsis. BMC Genomics 18, 538 (2017).

42. Schindler, U., Beckmann, H. & Cashmore, A. R. HAT3.1, a novel Arabidopsis homeodomain protein containing a conserved cysteine-rich region. Plant J. 4, 137–150 (1993).

43. Ueda, M. et al. Versatility of HDA19-deficiency in increasing the tolerance of Arabidopsis to different environmental stresses. Plant Signal. Behav. 13, e1475808 (2018).

44. Iuchi, S. et al. Regulation of drought tolerance by gene manipulation of 9-cis - epoxycarotenoid dioxygenase, a key enzyme in abscisic acid biosynthesis in Arabidopsis. Plant J. 27, 325–333 (2001).

45. Yang, S. U., Kim, H., Kim, R. J., Kim, J. & Suh, M. C. AP2 / DREB Transcription Factor RAP2. 4 Activates Cuticular Wax Biosynthesis in Arabidopsis Leaves Under Drought. Front. Plant Sci. 11, 895 (2020).

46. Varoquaux, N. et al. Transcriptomic analysis of field-droughted sorghum from seedling to maturity reveals biotic and metabolic responses. Proc. Natl. Acad. Sci. U. S. A. 116, 27124–27132 (2019).

47. Phillips, K. & Ludidi, N. Drought and exogenous abscisic acid alter hydrogen peroxide accumulation and differentially regulate the expression of two maize RD22-like genes. Sci. Rep. 7, 1–12 (2017).

48. Hernández-Sánchez, I. E., Maruri-López, I., Graether, S. P. & Jiménez-Bremont, J. F. In vivo evidence for homo- and heterodimeric interactions of Arabidopsis thaliana dehydrins AtCOR47, AtERD10, and AtRAB18. Sci. Rep. 7, 1–13 (2017).

49. Neha Pandey, Alok Ranjan, Poonam Pant, Rajiv K Tripathi, Farha Ateek, Haushilla P Pandey, U. V. P. and S. V. S. CAMTA 1 regulates drought responses in Arabidopsis thaliana. BMC Genomics 14, 216 (2013).

50. Seo, P. J. & Park, C. M. Cuticular wax biosynthesis as a way of inducing drought resistance. Plant Signal. Behav. 6, 1043–1045 (2011).

51. Yang, X. et al. Overexpression of an aquaporin gene EsPIP1;4 enhances abiotic stress tolerance and promotes flowering in Arabidopsis thaliana. Plant Physiol. Biochem. 193, 25–35 (2022).

52. Xu, Y. H. et al. Light-harvesting chlorophyll a/b-binding proteins are required for stomatal response to abscisic acid in Arabidopsis. J. Exp. Bot. 63, 1095–1106 (2012).

53. Dobin, A. et al. STAR: ultrafast universal RNA-seq aligner. Bioinformatics 29, 15–21 (2013).

54. Trapnell, C. et al. Differential gene and transcript expression analysis of RNA-seq experiments with TopHat and Cufflinks. Nat. Protoc. 7, 562 (2014).

55. Ernst, J. & Bar-Joseph, Z. STEM: A tool for the analysis of short time series gene expression data. BMC Bioinformatics 7, 191 (2006).

56. Saeed, A. I. et al. TM4: A free, open-source system for microarray data management and analysis. Biotechniques 34, 374–378 (2003).

57. Saleh, A., Alvarez-Venegas, R. & Avramova, Z. An efficient chromatin immunoprecipitation (ChIP) protocol for studying histone modifications in Arabidopsis plants. Nat. Protoc. 3, 1018–1025 (2008).

58. Li, H. & Durbin, R. Fast and accurate short read alignment with Burrows-Wheeler transform. Bioinformatics 25, 1754–1760 (2009).

59. Zhang, Y. et al. Model-based analysis of ChIP-Seq (MACS). Genome Biol. 9, R137 (2008).

60. Klein, H. & Sch, M. epigenomix — Epigenetic and gene transcription data normalization and integration with mixture models. (2017).

61. Mehdi, S. et al. MSI1 functions in a HDAC complex to fine-tune ABA signaling. Plant Cell 28, 42–54 (2016).

62. Zhang, X., Henriques, R., Lin, S., Niu, Q. & Chua, N. Agrobacterium -mediated transformation of Arabidopsis thaliana using the floral dip method. Nat. Protoc. 1, 1–6 (2006).

63. Jiang, Y., Liang, G. & Yu, D. Activated expression of WRKY57 confers drought tolerance in arabidopsis. Mol. Plant 5, 1375–1388 (2012).

