## Extended Figure for "SiHDA9 interacts with SiHAT3.1 and SiHDA19 to repress dehydration responses through H3K9 deacetylation in foxtail millet"

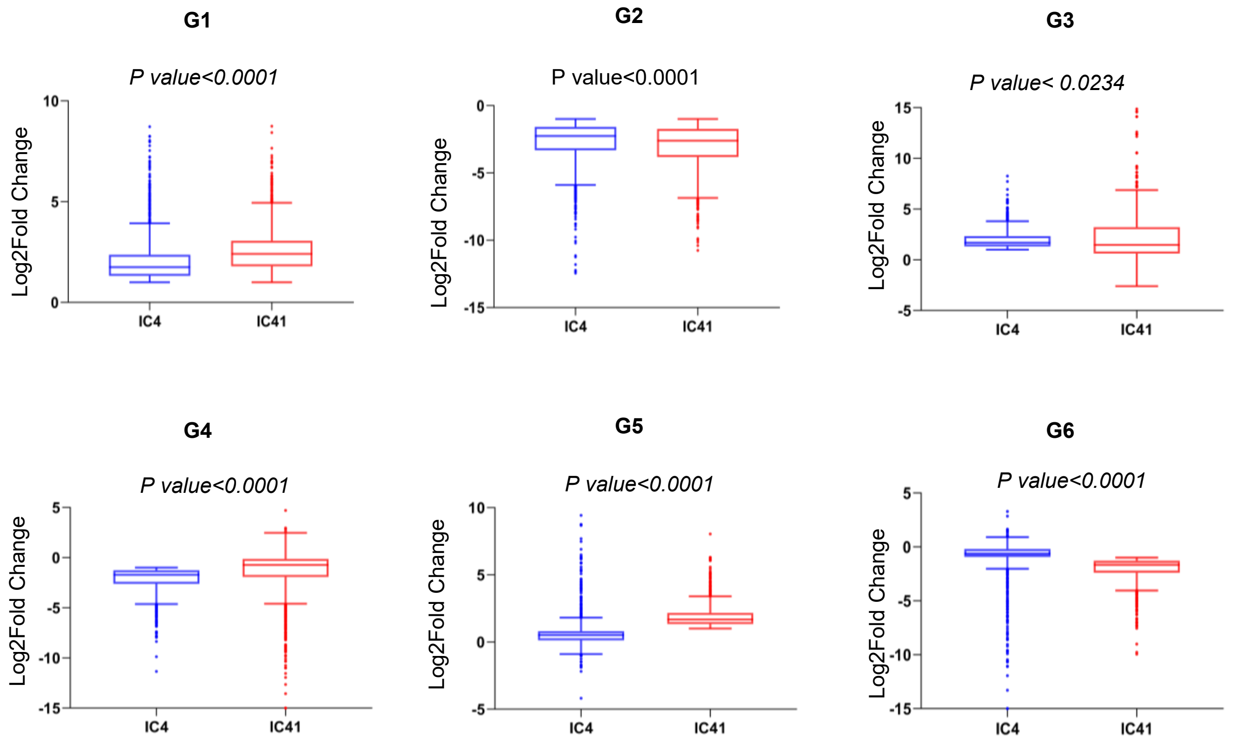


**Extended Data Fig. 1 |** Box plot showing the expression level (log2Fold change) of the genes belonging to the different groups between IC4 (drought tolerant) and IC41 (drought-sensitive) cultivars post 24hr PEG treatment. The G1-6 showing the groups of genes common with similar expression patterns **G1** IC4_IC41-UP (2418 genes) and **G2** IC4_IC41-DN (2177 genes), with unique expression patterns **G3** IC4_UP (1185 genes), **G4** IC4_DN (1245 genes), **G5** IC41_UP (2126 genes) and **G6** IC41_DN (919 genes). The *p-value* shows the significant level of gene expression between both cultivars. UP and DN stands for UP-regulated and Down-regulated genes, respectively.

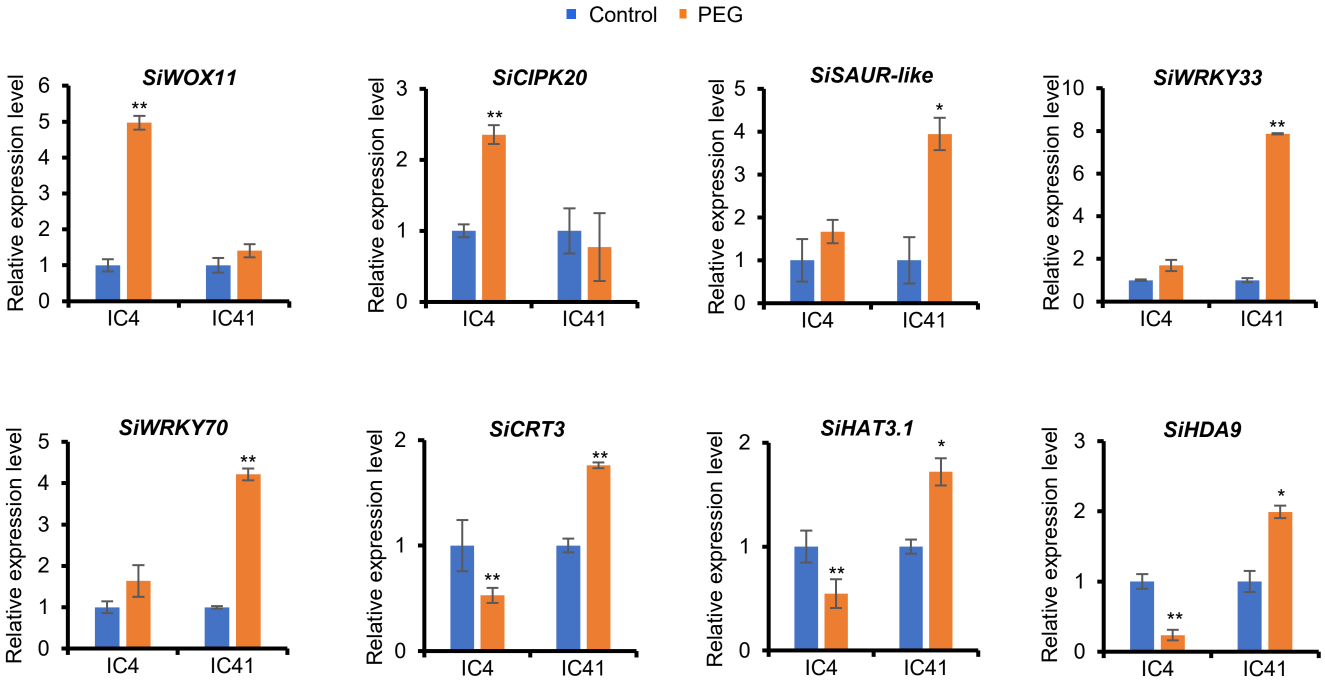


**Extended data Fig. 2 |** Validation of expression pattern of randomly selected DEGs of RNA-sequencing data by qRT-PCR after PEG (24hrs) treatment in IC4 and IC41 cultivars. Error bars represent standard error between three biological replicates. Asterisks (*) represent Student's t-test: * $P < 0.05$; ** $P < 0.01$.

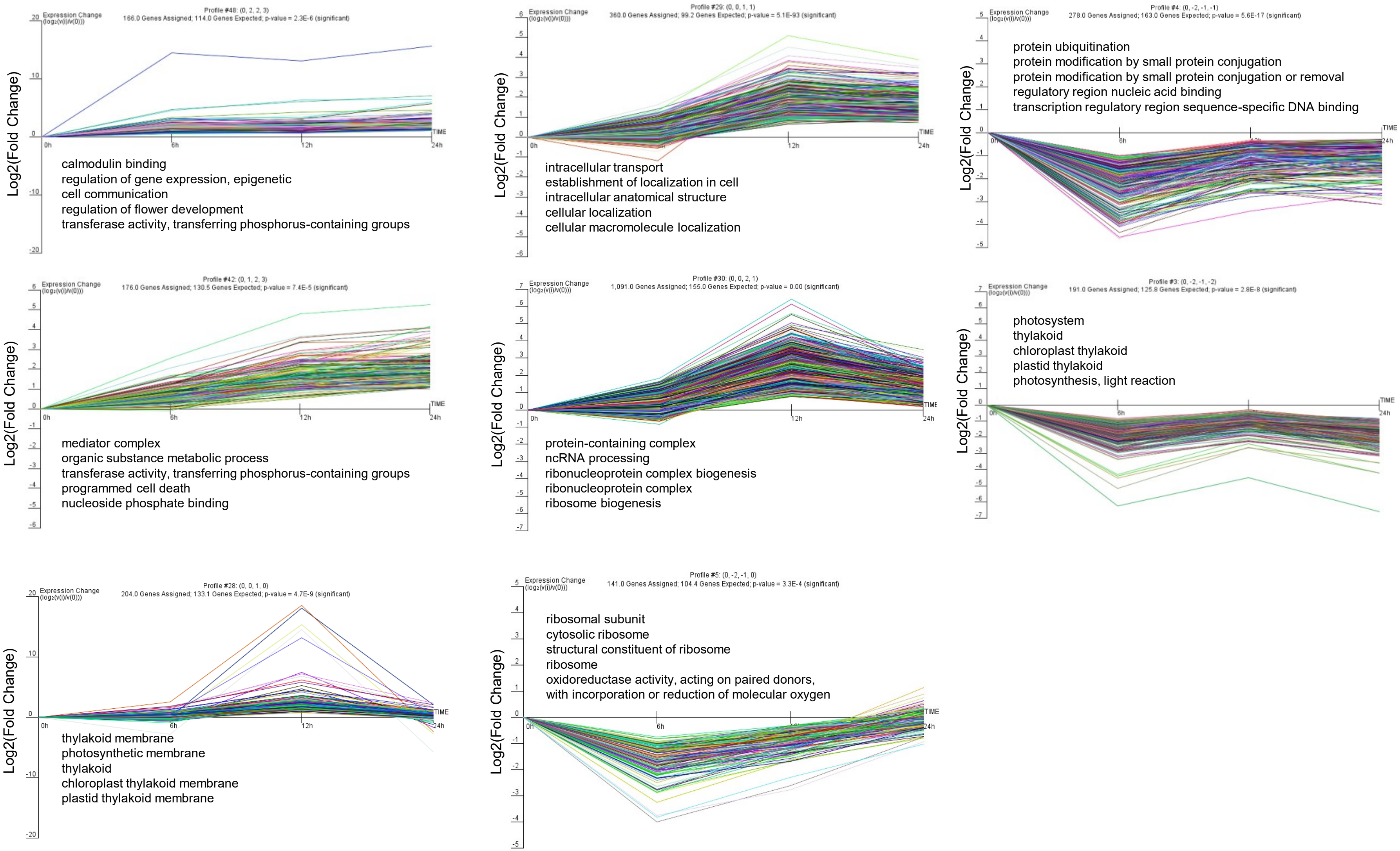


**Extended Data Fig. 3 | A cluster analysis revealed eight other main clusters in the PEG treatment time-course experiment in IC41 in comparison to the IC4 cultivar.** Clusters visualize the log2 (Fold change) expression dynamics after PEG-treatment. The five strongest enriched GO terms are shown for each cluster. For each time point (0, 6, 12, and 24hr), the average expression of IC41 was normalized with the average expression of IC4 (IC41/IC4) of RNA-sequencing data.

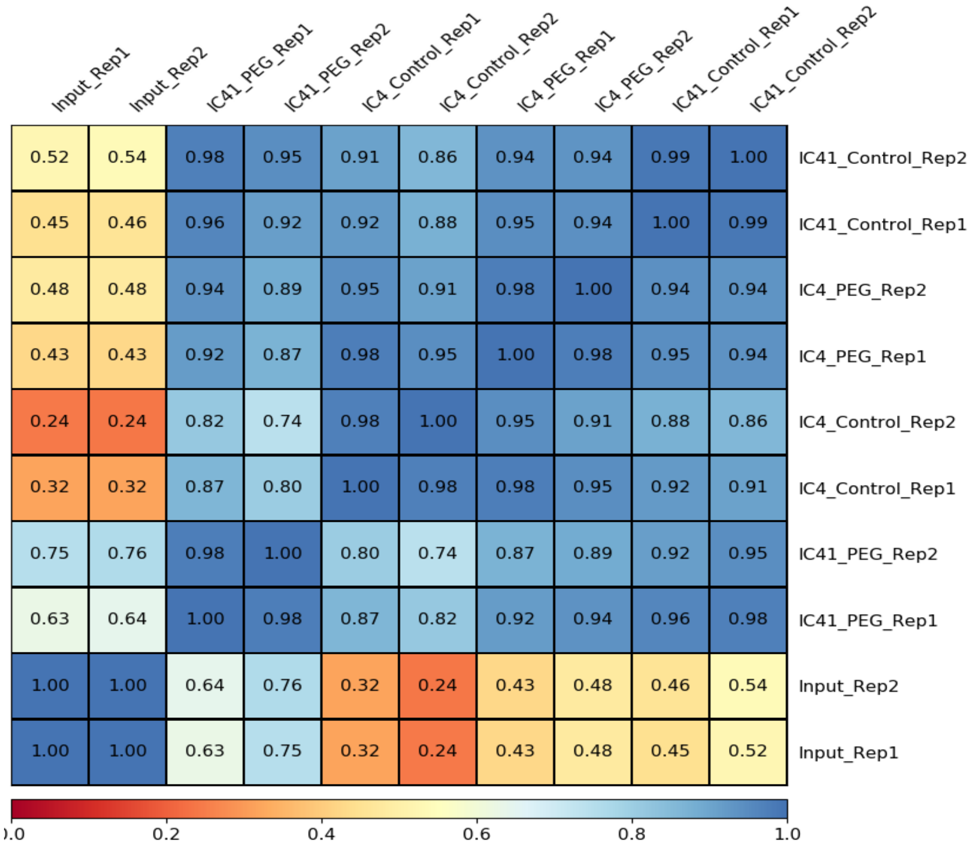


**Extended Data Fig. 4 |** A correlation matrix of ChIP (H3K9ac) samples of control (0hr) and PEG treatments (24hr) between biological replicates of IC4 and IC41 cultivars of foxtail millet. Values in the box show the correlation scores between samples.

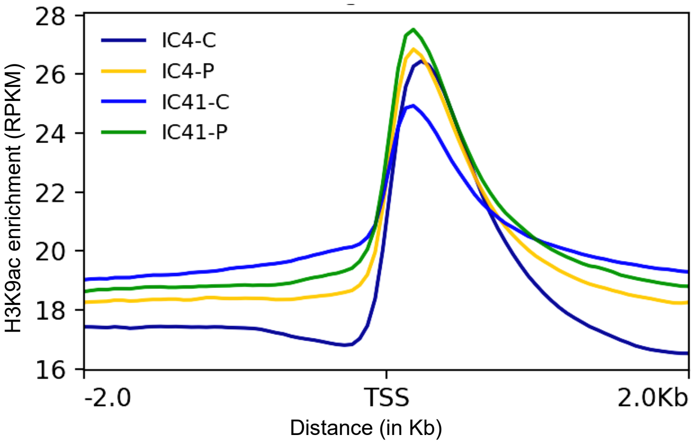


**Extended data Fig. 5 |** The enrichment of H3K9ac genic peaks around the transcription start site in all the samples. IC4 and IC41 are drought- tolerant and sensitive cultivars, respectively. C and P denote control and PEG-treated (24hr) samples, respectively. TSS is the transcription start site.

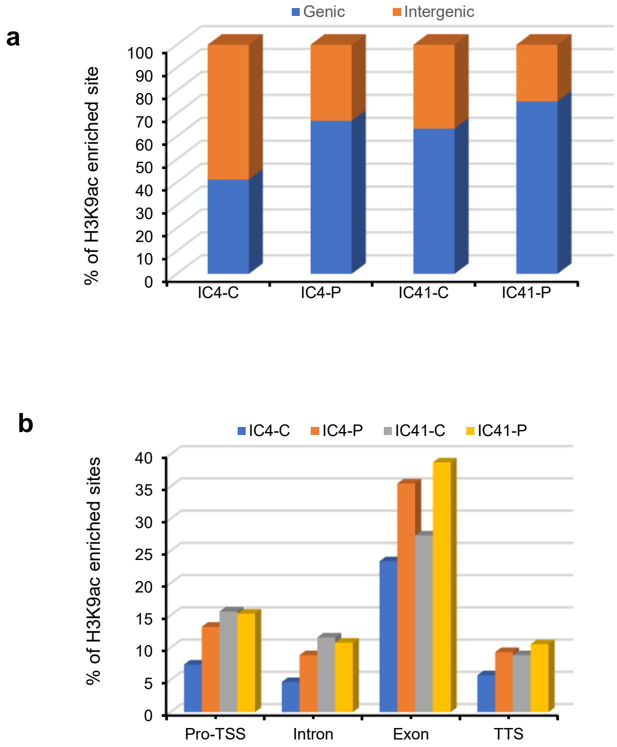


**Extended data Fig. 6 | The status of H3K9ac enriched peaks in control (IC4-C and IC41-C) and PEG-treated (IC4-P and IC41-P) samples**. **a)** Distribution of enriched peaks in genic and intergenic regions. **b)** Categorization of genic peaks according to their distribution on different regions of genes; Pro-TSS (Promoter-Transcription Start Site), Intron, Exon, and TTS (Transcription Termination Site) in IC4-C, IC4-P, IC41-C, IC41-P. C and P stand for control (0hr) and PEG-treated (24hr) samples, whereas IC4 and IC41 stand for dehydration-tolerant and sensitive cultivars, respectively.

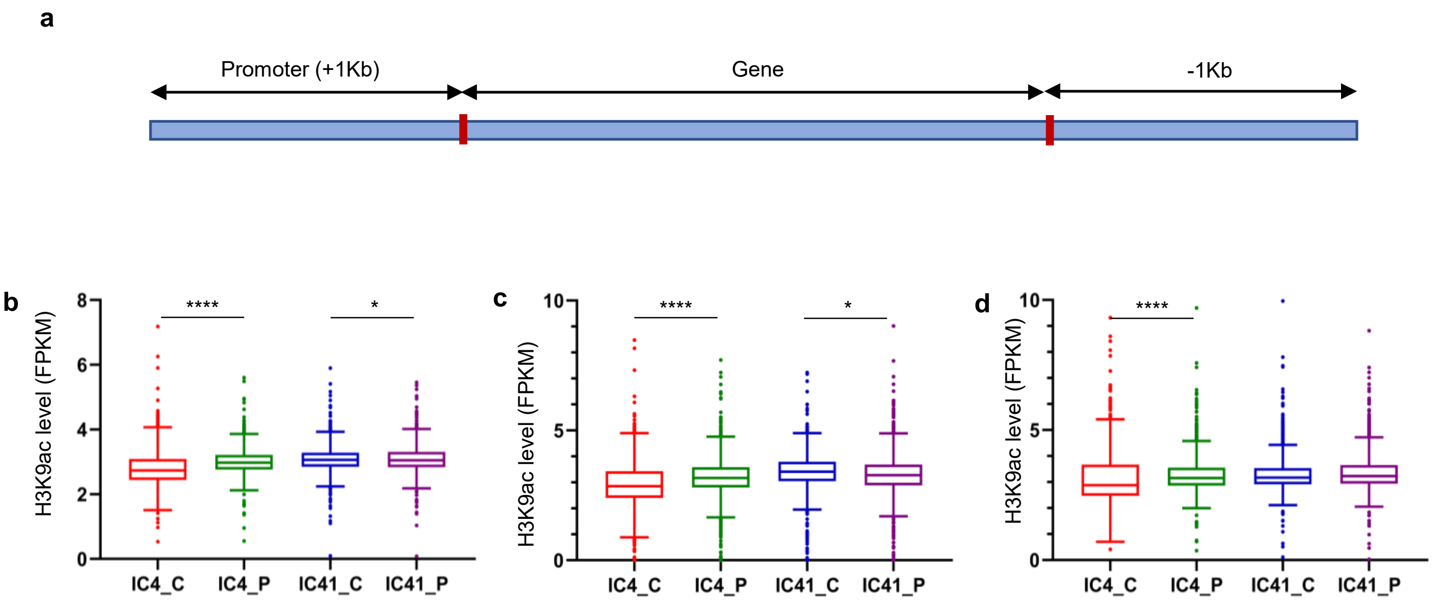


**Extended data Fig. 7 | Status of H3K9ac level on the responses to stress encoding unique DEGs (between control and 24hr PEG treated samples) found in the RNA-seq data of IC4 and IC41 cultivars. a)** Gene structure of a gene. The estimation of H3K9ac level (FPKM)) showed a enrichment (in IC4) and depletion (in IC41) of H3K9ac level on the **b)** gene with ± 1Kb and **c)** promoter (-1Kb) region of stress-responsive DEGs of IC4 and IC41. **d)** If H3K9ac was estimated only on the gene then it was significantly enriched in IC4 but unchanged in IC41. A total of 1445 stress-responsive unique DEGs (IC4-UP (476), IC4-DN (456), IC41-UP (652), and IC41-DN (447)) were included for the H3K9ac estimation. FPKM; fragments per kilobase per million mapped reads. Asterisks represented the student's t-test: * $P < 0.05$ and **** $P < 0.0001$.

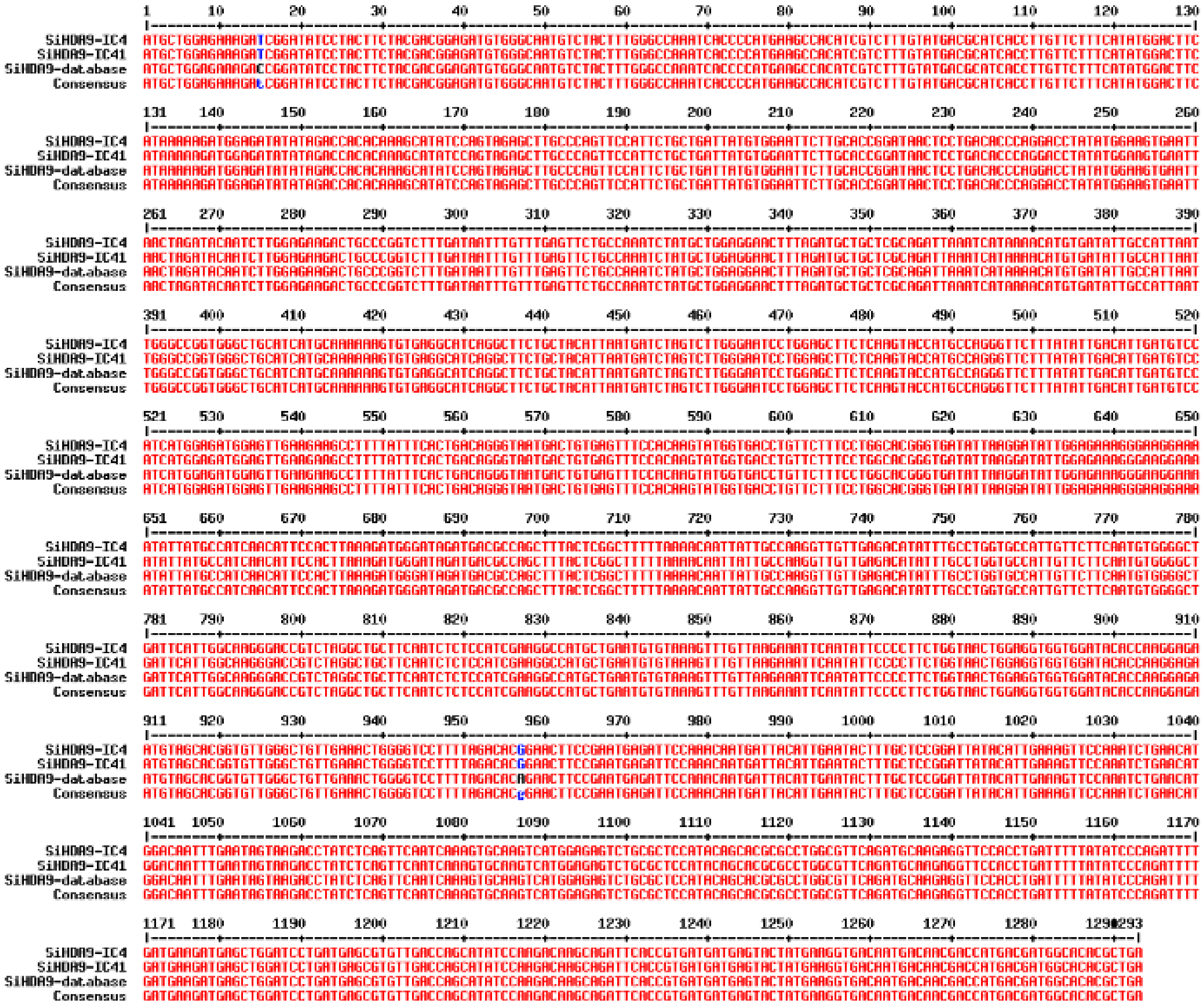

**Extended data Fig. 8 |** Sequence alignment showing no differences between coding sequence of *SiHDA9* of IC4 and IC41 cultivars of foxtail millet. However, two differences (from C to T at the 15th position and A to G at the 957th position) were found between our and the sequence in the database (phytozome).

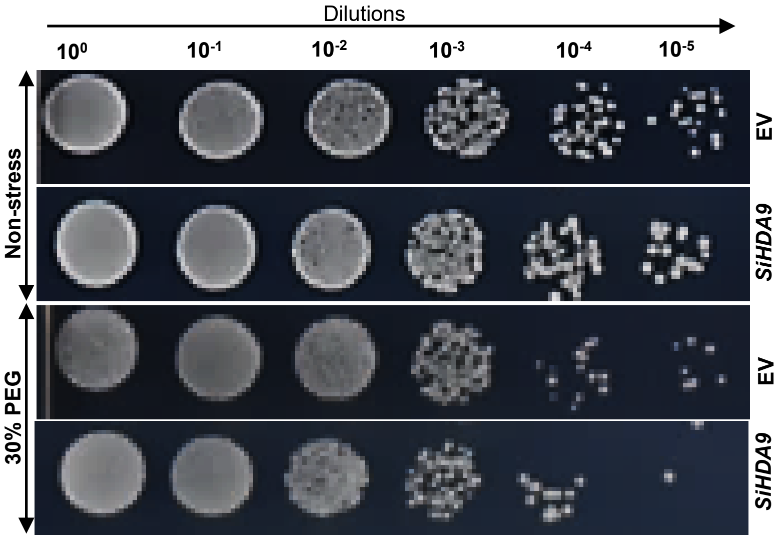


**Extended data Fig. 9 |** Stress tolerance assay for dehydration assay. *SiHDA9* overexpressed lines of yeast (*Saccharomyces cerevisiae*) are more susceptible upon the 30% PEG treatment respective to the control cells (with EV: empty vector), whereas no differences between them in the non-stress condition.

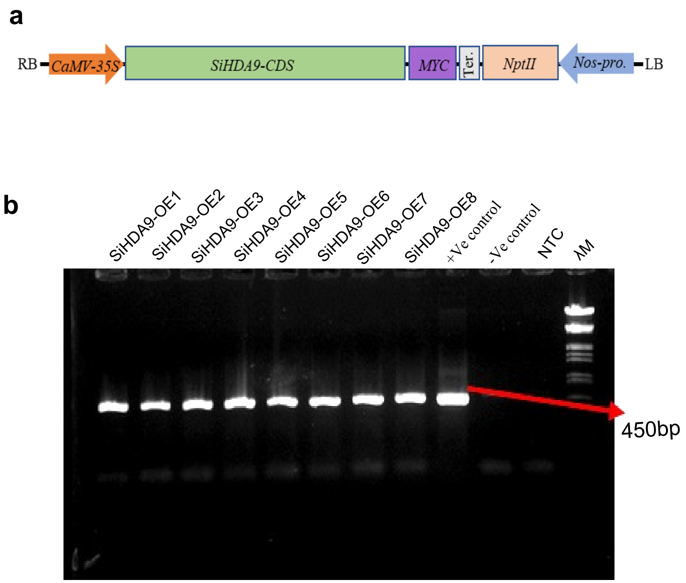


**Extended data Fig. 10 | Screening of SiHDA9 transgenic lines in Arabidopsis. a)** Schematic representation of *SiHDA9* construct used for Agrobacterium-mediated transformation in *Arabidopsis thaliana*. **b)** Gel image showing the PCR positive lines of *SiHDA9*. The PCR was performed using the *nptII (Neomycin phosphotransferase)* gene-specific primers.

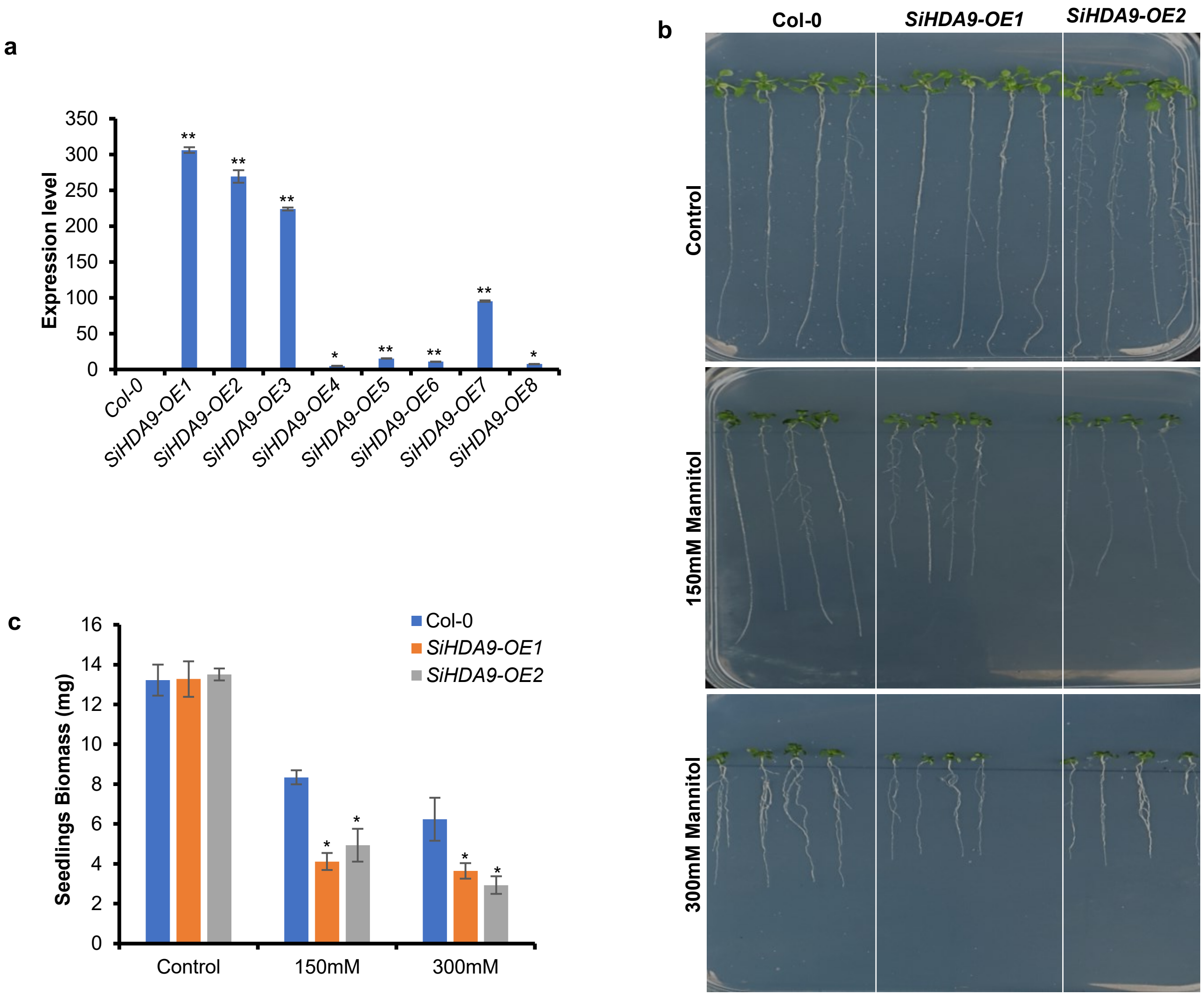


**Extended data Fig. 11 | *SiHDA9* overexpression lines showed more sensitivity upon mannitol treatment. a)** The expression analysis by RT-PCR showed the *SiHDA9* expression only in its overexpressing transgenic lines (T2) but not in Col-0 (wild-type). **b)** For the stress treatment assay, the T3-generation seeds of Col-0, *SiHDA9-OE1*, and *SiHDA9-OE2* were germinated on the 0.5MS-Agar plates for one week, and then seedlings were transferred on the 150mM and 300mM concentration containing 0.5MS-Agar plates and grow for another week before capture photographs. The results showed *SiHDA9* overexpressing plants were more sensitive to the dehydration stress (mimicked by mannitol) and suppressed their growth compared to the Col-0, with no significant differences between them in control (unstressed) conditions. **c)** The average biomass was significantly lower in both transgenic lines respective to the Col-0. The average of three plates (4 seedlings in each) was plotted. Error bars represent the standard error of three biological replicates. Asterisks represent the student's t-test: ** $P < 0.01$ and * $P < 0.05$.

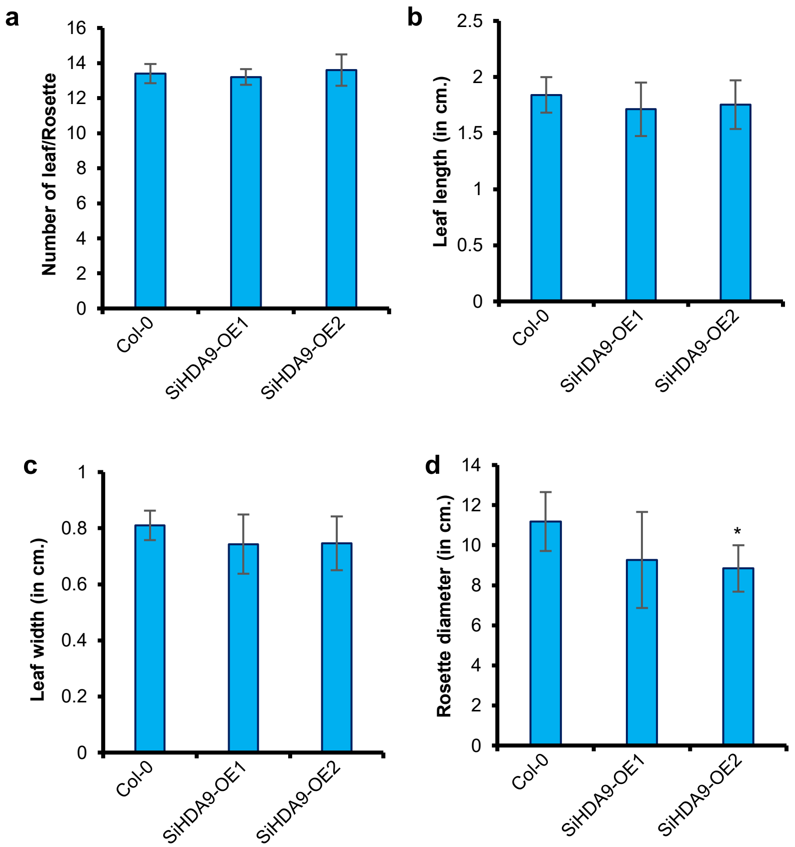


**Extended data Fig. 12 |** Phenotypic data at 30 DAS (Day After Sowing) showed no significant differences in **a)** leaf number, **b)** leaf length, and **c)** leaf width except **d)** rosette diameter between *SiHDA9* overexpression lines and Col-0. Error bars represent the standard error of five biological replicates. Asterisks represent the student's t-test: * $P < 0.05$.

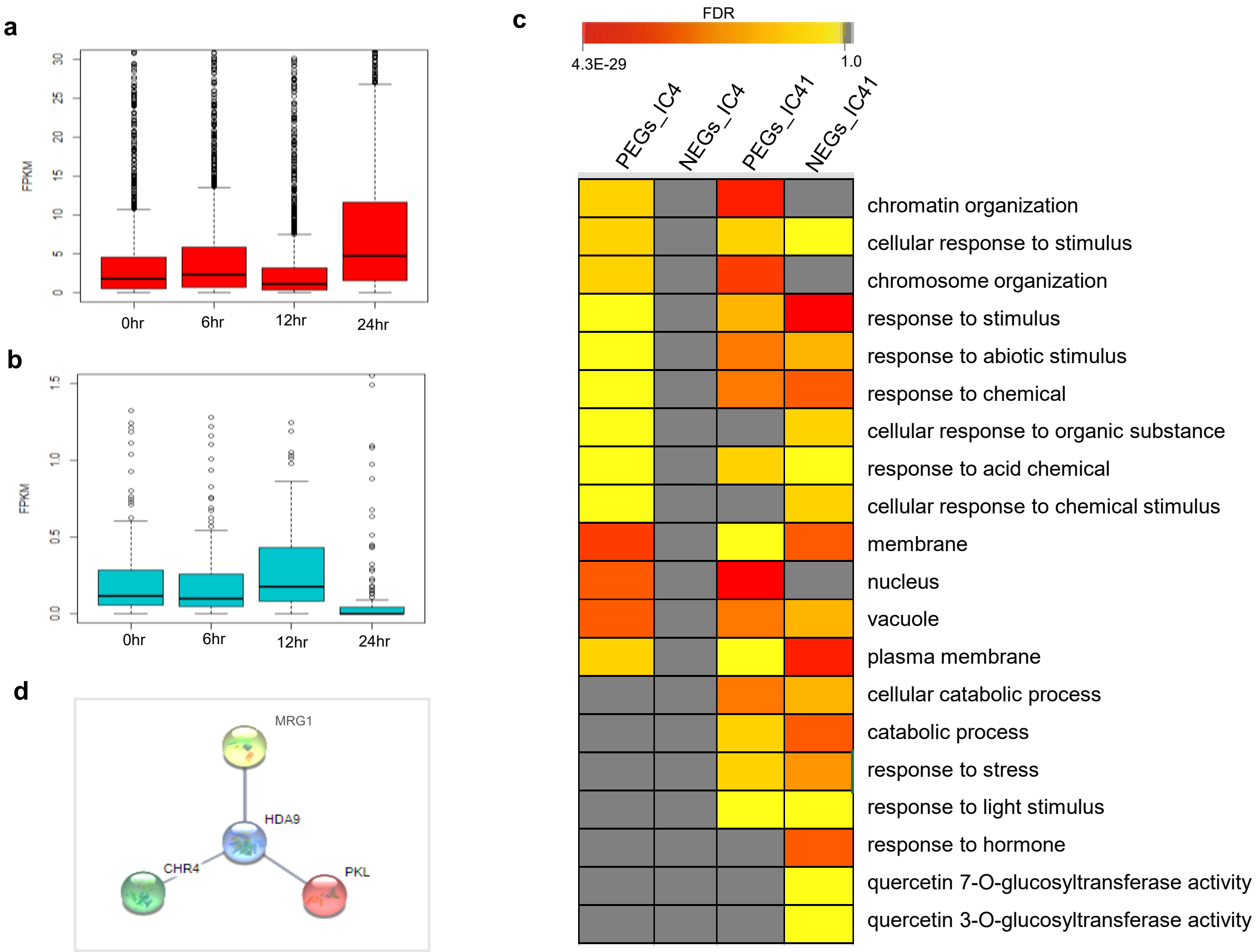


**Extended data Fig. 13 | Analysis of co-expression correlation of genes with *SiHDA9*.** Box plot showing the expression pattern of **a)** Positively & **b)** Negatively co-expressed genes at 0, 6, 12 & 24 hrs. in IC4 (tolerant cultivar). c) AgriGO analysis of positively co-expressed genes (PEGs) and negatively co-expressed genes (NEGs) showing enriched pathways in both IC4 and IC41 cultivars. d) String result showing the SiHDA9 interacting proteins in IC4 at confidence level 0.700.

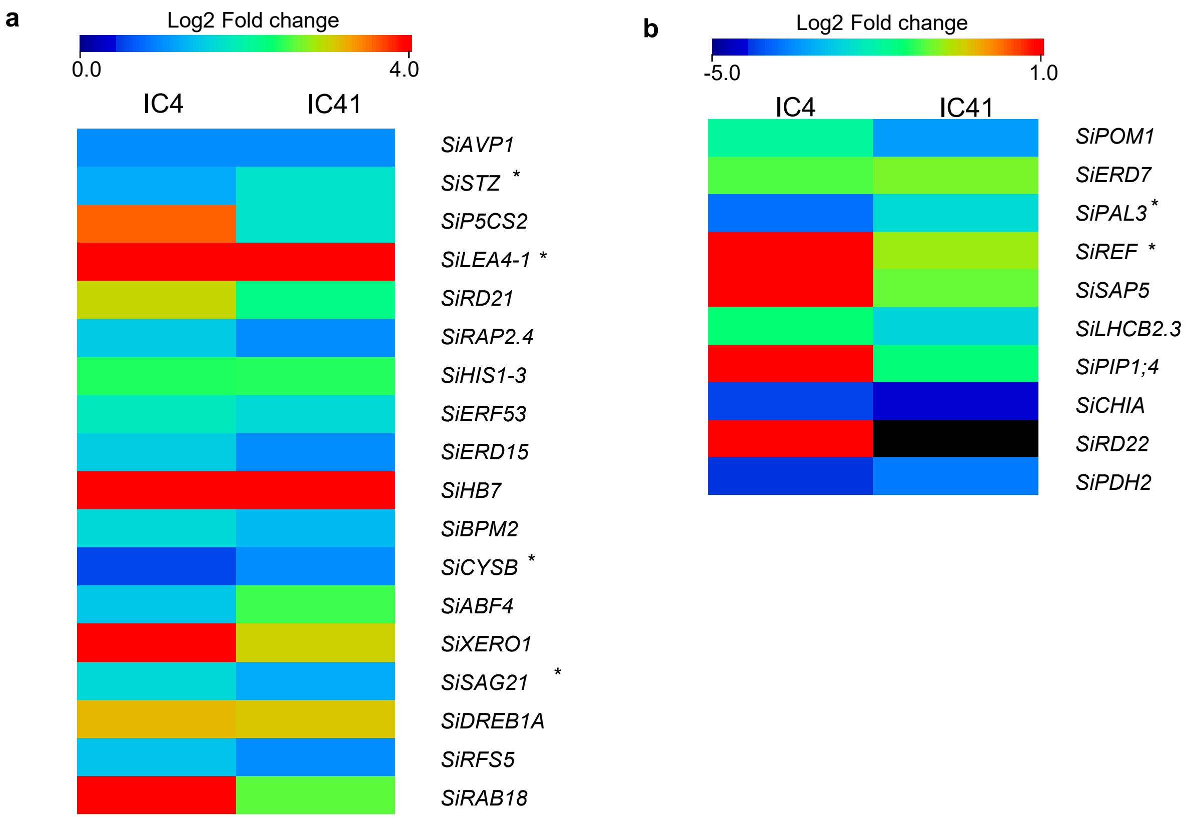


**Extended data Fig. 14 I** Heatmap showing the differences in expression level (between cultivars IC4 and IC41) of water deprivation pathway encoding genes found uniquely in **a)** IC4-cUP-gUP and b) **)** IC41-cDN-gDN post 24hr PEG treatments. All the genes possess HAT3.1 binding motif in their promoter except genes indicated by asterisks.

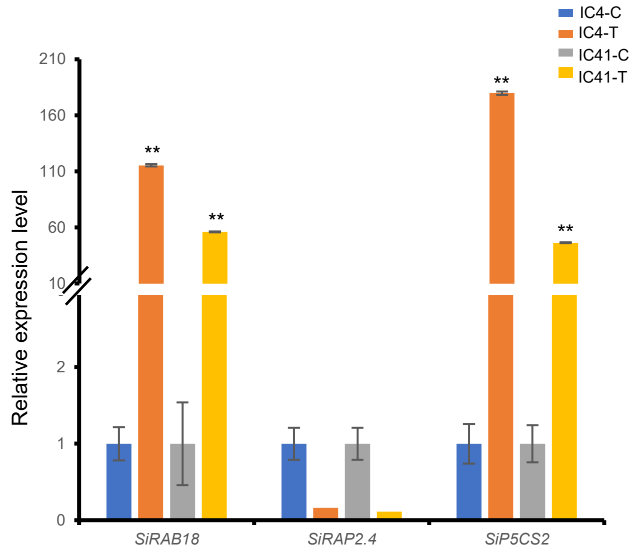


**Extended data Fig. 15 | Expression analysis of *SiRAB18*, *RAP2.4* and *SiP5CS2* after TSA (Trichostatin A) treatment (HDAC inhibitor) to the 21days old seedlings of dehydration tolerant and dehydration sensitive cultivar.** C and T in the sample names represent the control and TSA, respectively. Error bars represent the standard error between biological replicates. Asterisks represent the student's t-test: ** $P < 0.01$.
